# Evidence for abstract codes in parietal cortex guiding prospective working memory

**DOI:** 10.1101/2025.02.17.638684

**Authors:** Jongmin Lee, David De Vito, Jacob A. Miller, Derek Evan Nee

## Abstract

The recent past helps us predict and prepare for the near future. Such preparation relies on working memory (WM) which actively maintains and manipulates information providing a temporal bridge. Numerous studies have shown that recently presented visual stimuli can be decoded from fMRI signals in visual cortex (VC) and the intra-parietal sulcus (IPS) suggesting that these areas sustain the recent past. Yet, in many cases, concrete, sensory signals of past information must be transformed into the abstract codes to guide future cognition. However, this process remains poorly understood. Here, human participants used WM to maintain a separate spatial location in each hemifield wherein locations were embedded in a learned spatial sequence. On each trial, participants made a sequence-match decision to a probe and then updated their WM with the probe. The same abstract sequence guided judgments in each hemifield, allowing the separate detection of concrete spatial locations (hemifield-specific) and abstract sequence positions (hemifield-general), and also tracking of representations of the past (last location/position) and future (next location/position). Consistent with previous reports, concrete past locations held in WM could be decoded from VC and IPS. Moreover, in anticipation of the probe, representations shifted from past to future locations in both areas. Critically, we observed abstract coding of future sequence positions in the IPS whose magnitude related to speeded performance. These data suggest that the IPS sustains abstract codes to facilitate future preparation and reveal a transformation of the sensory past into abstract codes guiding future behavior.

**Significant Statement:** To act efficiently, we must use the recent past to prepare for what comes next. For this purpose, working memory (WM) is critical. Although substantial research has shown that WM retains recently presented sensory information, preparation for the future involves abstraction wherein shared meaning is aggregated while superfluous sensory details are discarded. The mechanisms underlying this process remain unclear. Analyzing functional MRI signals in the intraparietal sulcus (IPS), we found that distinct sensory states with shared predictive meaning were initially maintained in a sensory-like fashion, but over time, became aggregated indicative of abstraction. Abstraction was associated with behavioral efficiency highlighting its role in preparation. These findings reveal neural mechanisms supporting the transformation from past to future in WM.

## Introduction

Imagine performing instructed boxing combinations such as jab twice with the lead hand, hook with the other, then uppercut with the lead hand. To do so, the combination needs to be held in mind with each strike chaining into the next. This task requires working memory (WM), which refers to the maintenance and manipulation of information guiding ongoing behaviors (Baddeley, 1992; Curtis and D’Esposito, 2003; D’Esposito and Postle, 2015; Nee and D’Esposito, 2018). WM is central to much of higher-level cognition as many of our behaviors require extending the recent sensory past in order to properly act in the moment and near future.

Much of our use of WM is prospective in nature (Fuster, 1990; Rainer et al., 1999; Fuster, 2001; Nobre and Stokes, 2019) as the recent past allows us to prepare for the near future. Yet, most WM studies solely focus on how recent sensory content is maintained. Standard paradigms present participants with sensory information such as colors, orientations, spatial locations, or motion directions and then ask participants to recognize or reproduce the previously presented stimuli. A wealth of studies using such paradigms have demonstrated that high-fidelity stimulus-specific codes can be decoded from fMRI activation patterns in a distributed set of regions across visual and parietal cortices (Harrison and Tong, 2009; Serences et al., 2009; Christophel et al., 2012; Albers et al., 2013; Sprague and Serences, 2013; Sprague et al., 2014; Christophel et al., 2015; Ester et al., 2015; Bettencourt and Xu, 2016; Sprague et al., 2016; Lorenc et al., 2018; Rademaker et al., 2019; Li and Curtis, 2023). Findings such as these have given rise to the sensory recruitment hypothesis which states that WM is supported by the sustained maintenance of sensory information in those same sensory cortices involved in initial perception (Pasternak and Greenlee, 2005; Serences, 2016; Scimeca et al., 2018; Sreenivasan and D’Esposito, 2019).

While substantial sensory information is held in mind to support WM, much of our WM is guided by non-sensory codes. In particular, many of our prospective actions are guided by more abstract codes. For example, we can apply the same boxing combination described above when in an orthodox (left foot/hand forward) or southpaw stance (right foot/hand forward) such that the hand performing each strike is reversed as a function of stance. Abstraction is a process by which shared features are aggregated or compressed together (Ho et al., 2019; Courellis et al., 2024). This process affords generalization which facilitates adaptive behavior and knowledge transfer across contexts (Badre et al., 2010; Collins and Frank, 2013; Badre, 2024). Despite the importance of generalization to WM and cognition, few studies have examined abstract coding in support of WM. Some evidence has demonstrated that concrete sensory information of the past can be recoded into a more abstract format such as translating a motion cloud to a line (Kwak and Curtis, 2022; Duan and Curtis, 2024). Moreover, some recent studies have demonstrated that when forthcoming actions can be anticipated, prospective action codes are maintained (van Ede et al., 2019; Boettcher et al., 2021; Henderson et al., 2022; Nasrawi and van Ede, 2022; Shushruth et al., 2022). However, there is a critical gap in our understanding of the abstract representations that reside in-between sensation and action which prospectively and flexibly guide WM for the future.

Abstract coding increases with synaptic distance from primary cortices (Mesulam, 1998; Brincat et al., 2018). For visuo-spatial processing, the intra-parietal sulcus (IPS) is downstream of visual cortical areas of the occipital cortex (e.g., V1-V4; VC). Although in numerous cases, visual WM is decoded similarly in VC and IPS (Christophel et al., 2012; Sprague and Serences, 2013; Sprague et al., 2014, 2016) some work has demonstrated that IPS WM representations are more robust to distraction (Bettencourt and Xu, 2016; Lorenc et al., 2018; Rademaker et al., 2019). It has been speculated that abstraction results in a stable, low-dimensional coding format that is distractor resistant (Murray et al., 2017; Rademaker et al., 2019). Hence, we reasoned that abstract codes that facilitate prospective uses of WM would be relatively more prominent in the IPS than VC.

The current study used fMRI to investigate the mechanisms underlying the use of the recent past to prepare for the future. Human participants performed a sequence-matching task which probed whether a presented spatial location followed the last spatial location in a predefined sequence (**Figure 1**). The sequence (**Figure 1a**) was abstract in that it could be generalized to apply to spatial locations in either hemifield. That is, we define abstraction as the ability to generalize over contexts or instances (Badre and D’Esposito, 2007; Nee and D’Esposito, 2016; Bernardi et al., 2020; Badre et al., 2021; Kwak and Curtis, 2022; Courellis et al., 2024; Duan and Curtis, 2024). Moreover, the use of a sequence afforded the use of the recent past (most recently presented location) to predict and prepare for the near future (next location in the sequence). Through this task, we examined concrete (spatial location), abstract (sequence position), retrospective (previous location/position), and prospective (next location/position) coding in visual and parietal cortices. We predicted that both VC and IPS would show initial evidence of concrete, retrospective coding consistent with numerous past works. However, we anticipated the emergence of abstract and prospective coding in the IPS consistent with a top-down code to guide future expectations. Such a result would provide evidence for the use of abstract codes to guide prospective WM.

**Figure 1.**
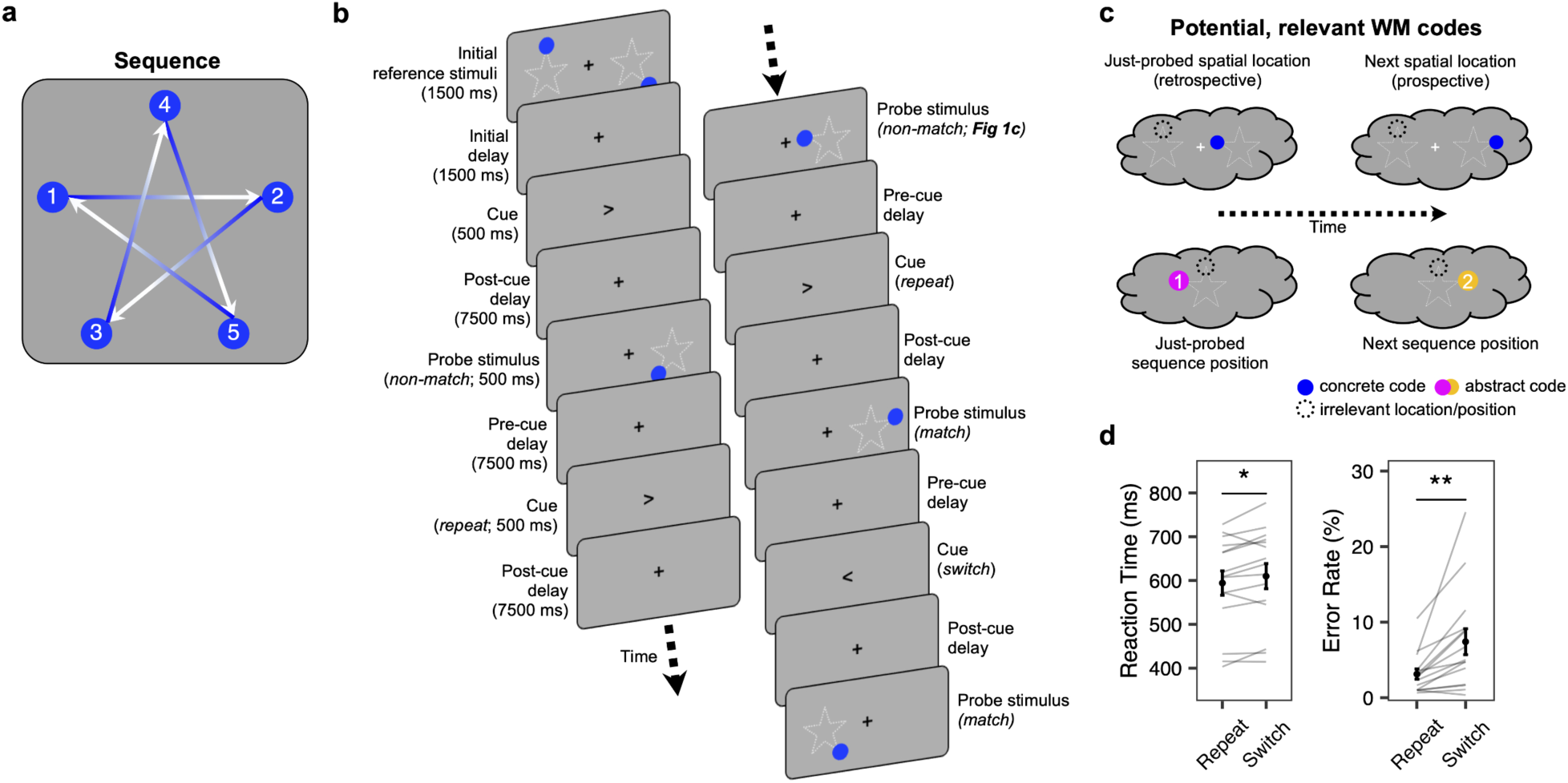
Procedures and behavioral performance. **(a)** The sequence consisted of five points forming a star. Arrows indicate the sequence ordering such that position 2 followed position 1, position 3 followed position 2, etc. Position 5 transitioned back to position 1 such that the sequence was circular. **(b)** Task procedure. Each trial consisted of a pre-cue delay, a cue, and a post-cue delay. At the start of each run, initial spatial locations were presented in the left and right visual hemifields. After an initial delay, a cue indicated the task-relevant visual hemifield wherein a forthcoming probe would be presented following a post-cue delay. Participants determined whether the spatial location of the probe followed the spatial location of the last item that appeared in the same hemifield. Half of the probes were sequence matches (e.g., position 2 following position 1) while half were non-matches (e.g., position 3 following position 1). The dotted lines indicating the star trace are used for illustrative purposes and did not appear in the actual task. **(c)** The potential transformation of relevant WM codes over time. After the probe stimulus presentation, the spatial location of the just-probed stimulus (blue; concrete, retrospective code; hemifield-specific) and its sequence position (pink; abstract, retrospective code; hemifield-general) could be maintained in WM. To optimally prepare for the future, these hypothetical codes would be transformed over time to the next location(blue)/position(orange). Irrelevant (uncued) representations are included for completeness (dashed lines), but are not a main focus here. **(d)** Mean reaction time and error rates. Behavioral performance was better when the cue indicated the same visual hemifield as the previous trial (‘Repeat’) relative to when the cue indicated the other visual hemifield (‘Switch’). Each line on the bar graph represents each participant. Bold dots indicate the mean across participants with error bars representing ±1 standard error of mean. *p < 0.05, **p < 0.01.

## Materials and Methods

### Participants

22 participants were recruited from the Tallahassee area for this experiment. All participants underwent an initial screening process to ensure that they understood and could adequately perform the task. Informed consent was obtained according to the guidelines set forth by the Office for Human Subjects Protection and Florida State University Institutional Review Board. For the experiment, a total of three scanning sessions were planned. 2 participants were unable to tolerate the scanner environment, and 1 participant was lost due to a scheduling conflict. An additional 4 participants were removed from the study following the first scanning session due to insufficient data quality (e.g., 2 for poor task accuracy, 2 for excessive head motion). These data were excluded from further analyses. The remaining 15 participants (age range 18-23; mean age: 20.1; 10 female) were analyzed for this report. 1 participant did not complete the last scanning session due to a technical issue with the remaining participants completing all three sessions.

### fMRI experimental design

In the fMRI scanner, participants performed a sequence working memory task. In the task, participants were required to maintain two spatial locations, one in each visual hemifield, each of which were drawn from a five-location sequence tracing a star (**Figure 1**). These stimuli were used as references for the probes. On each trial, a cue was presented at the center of the screen for 500 ms, pointing to either the left or right visual hemifield, indicating that an upcoming probe would appear in that respective visual hemifield (task-relevant visual hemifield). For all but the first cue, cues were preceded (pre-cue delay) and followed (post-cue delay) by 7500 ms of fixation. After the post-cue delay, a probe was presented in the task-relevant visual hemifield for 500 ms. Participants were asked to determine whether the spatial location of the currently presented probe sequentially followed the reference (i.e., the spatial location of the last stimulus presented in the same visual hemifield). The spatial location of the probe then was used as the reference for the next probe in the same visual hemifield. Thus, the reference in the probed hemifield was updated in each trial.

Half of the probes followed the reference in the sequence thereby requiring a “match” response. “Non-match” probes were pseudo-randomly and equally drawn from the three spatial locations remaining after excluding the sequence “match” and the reference (i.e., there were no stimulus repeats). Trials were equally distributed to cue the left and right visual hemifields and to require even numbers of switches and repeats of the visual hemifield of the preceding trial. Each run began with the presentation of two location stimuli for 1500 ms which served as the initial reference locations. A delay of 2500 ms separated the initial stimuli from the first cue. During the experiment, participants were asked to fixate their gaze on a fixation cross the center of the screen and not move their eyes. Adherence to these instructions was monitored by eye tracking. 14 out of 15 participants completed three scanning sessions, while one participant completed only two scanning sessions due to a technical issue. Each scanning session was carried out on separate days. Each scanning session consisted of 4 experimental runs and 50 trials per experimental run. There was a short break between experimental runs.

### Stimuli and apparatus

The experiment was programmed with E-Prime software version 2.0 (Psychology Software Tools, Inc., Pittsburgh, PA). The probe stimulus was a blue circle with a diameter of ∼2°. Stimuli were presented between 5.5° and 15.25° of visual angle left and right and between 3.32° and 4.68° of visual angle above and below a central fixation point.

### MRI acquisition

MRI data were collected on a Siemens 3T Prisma with a 32-channel head coil. The participant observed visual stimuli in the scanner using a mirror attached to the head coil, which reflected the visual stimuli onto a projection screen. Responses were recorded using a 4-button MR-compatible button box (Current Designs, Inc.), with participants using the index finger of each hand to input their responses.

Functional imaging data were obtained using an EPI sequence provided by the Center for Magnetic Resonance Research (CMRR) at the University of Minnesota: voxel dimensions of 2 × 2 × 2 mm, TR = 2000 ms, TE = 33.8 ms, flip angle = 45°, FOV = 192 mm^2^, multi-band factor = 4. Prior to each functional scan, four dummy scans were conducted to ensure image stabilization. Phase and magnitude images at the same resolution of the functional images were acquired to assess and correct for magnetic field inhomogeneity. Additionally, a high-resolution T1-weighted MPRAGE image was collected for spatial normalization (384 × 384 ×256 matrix of 0.667 mm3 isotropic voxels; TR =1840 ms; echo time = 2.9 ms; flip angle = 9°).

### MRI data preprocessing

Preprocessing on image data was done in SPM12 (https://www.fil.ion.ucl.ac.uk/spm/) unless otherwise specified. DICOM format was converted into NIfTI format. The origin of all images in each participant was manually adjusted to the anterior commissure. Functional image data underwent spike correction to mitigate the influence of artifacts using AFNI’s 3dDespike routine (http://afni.nimh.nih.gov/afni), as well as slice timing correction for each run, head motion correction via a six-parameter rigid body transformation, and unwarping and correction of motion-by-susceptibility distortions using the FieldMap toolbox (Andersson et al., 2001). Based on the realignment parameters, linear, squared, differential, and squared differential movement parameters were calculated (24 in total; Lund et al., 2005). These parameters were regressed out of the functional images and the resultant residuals were high-pass filtered at 1/128 Hz. These data were used for all described analyses.

Structural image data were coregistered to the functional image data. Then, segmentation was performed on structural image data to acquire gray and white-matter probability maps (Ashburner and Friston, 1997) from which spatial normalization warping to and from the MNI template was calculated.

### ROI selection

We selected the left and right intra-parietal cortex (IPS) along with the left and right visual cortex (VC) as the regions of interest (ROIs) given their role in maintaining working memory (Harrison and Tong, 2009; Serences et al., 2009; Ester et al., 2015; Bettencourt and Xu, 2016; Lorenc et al., 2018; Rademaker et al., 2019). ROI masks were created using a probabilistic atlas (Wang et al., 2015) with a 90% probability threshold. VC ROIs were created by combining V1v, V1d, V2v, V2d, V3v, V3d, and hV4. IPS ROIs were created by combining IPS0-5. These ROIs were warped to native space for each individual.

### Multivariate fMRI analyses

The BrainIAK toolbox implemented in Python was used to conduct decoding analyses (Kumar et al., 2020). All decoding analyses were performed on preprocessed BOLD responses from the ROIs of each participant. To appropriately balance factors for classification, consecutive pairs of runs were concatenated. The concatenated BOLD data of each ROI were temporally z-scored to ensure that voxel activities were scaled to the same range. Only correct trials were analyzed. After removing error trials, we equalized the number of trials of each spatial location in each run to match the smallest count among them by randomly sampling the larger number of trials of each spatial location resulting in ∼8 trials per concatenated run per location on average. Such random sampling and cross-validation were repeated 1000 times.

To train and test a classifier (L2-regularized, multinomial logistic regression), we implemented a leave-one-pair-out cross-validation procedure. The decoding outputs of each participant was calculated as the average decoding outputs across all iterations. During cross-validation iteration, we applied univariate feature selection using an ANOVA (f-classif function in the scikit-learn Python library (Pedregosa, 2011) with a threshold alpha level of 0.05 on each training data. The features selected based on the training data were used on the testing data. Since this procedure resulted in a variable number of features as a function of ROI and time point, we repeated the analyses using a fixed number of features (300, which approximated the lower bound of the number of features selected by the above procedure) which produced the same patterns of results as reported in the main text (**Figure S6**) indicating that the results are robust to the specifics of feature selection. The L2 penalty was the value of 1, which was used to train the classifier. For the decoding analysis, we excluded the initial trial of each run as the timing and visual presentation of this trial differed from the others.

### Multivariate fMRI analyses: temporal generalization decoding analysis

Temporal generalization decoding analysis (King and Dehaene, 2014) was conducted to assess transformations of representations over time. Classifiers were trained to classify 10 distinct categories representing the spatial locations of stimuli (five locations in each hemifield) based on the average fMRI activation patterns over the last two time points of the post-cue delay period (12 and 14 seconds post-stimulus onset). Then, the trained classifier was tested to classify each TR of the independent testing data (8 TRs in total; from 0 to 14 s from the onset of the probe). For repeat trials, we assigned the same category label to 8 TRs of the pre-cue and post-cue delays, and the category of the label corresponded to the probe of the preceding trial. For switch trials, the 4 TRs of the pre-cue delay were labeled in the same way. However, the 4 TRs of the post-cue delay were labeled with the last probe presented on the cued visual hemifield. In other words, the pre-cue delay was always labeled with the last probe while the post-cue delay was always labeled with the reference for the upcoming probe.

Training and testing were performed separately for each of the four ROIs (left and right IPS and VC). Given similarity in the results in each hemifield, left and right hemifields were averaged together to summarize a given brain area (e.g., left and right IPS results were averaged to characterize the IPS).

The temporal generalization decoding analysis produced classifier evidence corresponding to each of the 10 spatial locations at each time point. We specifically tracked evidence for the retrospective-relevant (‘Retro-rel’), retrospective-irrelevant (‘Retro-irrel’), prospective-relevant (‘Pro-rel’), and prospective-irrelevant (‘Pro-irrel’) locations (**Figure 1c**). During the pre-cue delay, the just-probed spatial location was categorized as ‘Retro-rel’, and the last probed spatial location in the uncued hemifield was categorized as ‘Retro-irrel’. During the post-cue delay of the repeat condition, the next spatial locations of ‘Retro-rel’ and ‘Retro-irrel’ in the sequence were categorized as ‘Pro-rel’ and ‘Pro-irrel’, respectively. In the switch condition, the classifier evidence corresponding to the next spatial locations of ‘Retro-irrel’ and ‘Retro-rel’ in the sequence were categorized as ‘Pro-rel’ and ‘Pro-irrel’, respectively. For each time point, classifier evidence for each of the four categories of interest were averaged across trials for each subject and submitted to statistical analysis.

Prior work has suggested that representations during WM and perception are distinguishable (Rademaker et al., 2019; Lorenc et al., 2020). Hence, the analyses above focused on training the very end of the post-cue delay to maximize separation from sensory signals and hone in on mnemonic codes. For completeness, we repeated the procedures above using the last two time points of the pre-cue delay (4 and 6 seconds post-stimulus onset) to train classifiers. This timing corresponds to the peak of hemodynamic signals following a stimulus allowing the ability to determine the extent to which the results reflecting more sensory-like signals show the same patterns. These data are reported in Figure S4a.

### Multivariate fMRI analyses: cross-visual hemifield decoding analysis

Cross-visual hemifield decoding analysis was performed by training a classifier on the five sequence positions presented on the one visual hemifield (e.g., the left (right) visual hemifield) and testing the classifier on the five sequence positions presented on the other visual hemifield (e.g., the right (left) visual hemifield). Decoding was conducted in a time-resolved manner, involving training and testing the classifier within the same TR. As we did in the temporal generalization analysis, the label at each TR during the pre-cue delay was based on the just-probed stimulus, while the label at each TR during the post-cue delay was based on the last probed stimulus in the cued hemifield.

### Representational similarity analysis

We conducted a representational similarity analysis (RSA) (Kriegeskorte et al., 2008) on the preprocessed BOLD data to examine the transformation of the neural representation of WM across time points. To setup the data for RSA, consecutive pairs of runs were concatenated in the same way as in the decoding analyses. Then, data were split into training and testing sets by using a leave-one-pair-out-cross-validation procedure. For each iteration of the cross-validation, univariate feature selection was performed using an ANOVA with a threshold alpha level of 0.05 on the training data (note similar patterns of results were obtained by fixing the number of features at 300, **Figure S6**). The selected features (voxels) were then z-scored as a collection, followed by z-scoring within each feature. Based on these normalized features on the training data, we selected samples corresponding to the 10 spatial locations at the last two time points of the post-cue delay (12 and 14 seconds post-stimulus onset). Then, we calculated the average value of the selected samples for each spatial location to form the training representations (10 spatial locations). For the testing data, we used the same features selected on the training data and applied z-score normalization in the same manner as was done for the training data.

Then, we selected samples corresponding to the 10 spatial locations for each time point. For each time point, we averaged the patterns of activation for each of the 10 spatial locations, respectively, to form the testing representations. Then, for each time point, we calculated the correlation coefficient among the 10 training representations and the 10 testing representations. In other words, eight RSA matrices were formed, one per time point, showing the representational similarity between the 10 spatial locations of the training set and the 10 spatial locations in the testing set. Each RSA matrix was submitted to model-based regression analysis (see below) for each iteration of the cross-validation.

As we did for the temporal generalization decoding above, we repeated these procedures using pre-cue delay (4 and 6 seconds) trained data for completeness (**Figures S4b and S4c**).

### Representational similarity analysis: model-based regression

We performed multiple linear regression (Eq.1), using the vectorized correlation coefficient RSA matrices described above as the dependent variable (r) and a set of vectorized model RSA matrices as regressors. This analysis estimated the unique contribution of each model RSA matrix to the correlation coefficient RSA matrix. Below describes five model RSA matrices with details:

(1) A matrix representing hemifield spatial attention (‘SA’) reflecting similarity among stimuli presented on the same visual hemifield, but dissimilarity among stimuli presented on the opposite visual hemifields. Similarity was assigned a regressor value of 1 and dissimilarity was assigned a regressor value of -1.
(2) A matrix representing the spatial location of the reference (‘Concrete Retro’). Here, distance-dependent similarity was accounted for by computing the distance in visual angle among locations in the same hemifield. The distance in visual angle between two locations was scaled by the maximum distance among any two locations such that a location with itself received a value of 1 and the maximally distant two locations received a value of -1. Between hemifield locations were assigned a value of 0 as these dissimilarities are already captured by the SA regressor. Note that since we assume that training patterns reflect the prospective location, retrospective regressors are shifted back one location in the sequence (e.g., a trained pattern of L2 would be expected to appear as L1 early in the trial).
(3) A matrix representing the location following the reference in the sequence (‘Concrete Pro’). In this case, since we assume that training patterns reflect the prospective location, this matrix resembles the identity matrix along with distance-dependent similarity as above.
(4) A matrix representing the abstract sequence position for the reference stimulus (‘Abstract Retro’), wherein the values of the second model RSA matrix (‘Concrete Retro’) were flipped in the horizontal direction. By doing so, we modeled cross-hemifield generalizability of the abstract sequence position of the past. Note that the ideal matrix corresponding to a retrospective abstract sequence position would be the summation of the ‘Concrete Retro’ and ‘Abstract Retro’ matrices as described. However, such a matrix would induce collinearity with the ‘Concrete Retro’ matrix. Hence, the ‘Abstract Retro’ matrix as described captures additional variance attributed to cross-hemifield generalization over-and-above within-hemifield distance-dependent similarity.
(5) A matrix representing the abstract sequence position for the next location in the sequence (‘Abstract Pro’), wherein the values of the third model RSA matrix (‘Concrete Pro’) were flipped in the horizontal direction. By doing so, we modeled the cross-hemifield generalizability of the abstract sequence position of the upcoming future. As with the ‘Abstract Retro’ matrix, this matrix captures additional variance attributed to cross-hemifield generalization over-and-above within-hemifield distance-dependent similarity.

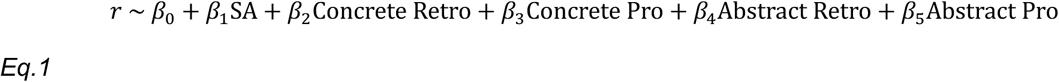

For each iteration of the cross-validation procedure, we estimated beta weights for each time point using the equation (Eq. 1). The beta weights obtained from the model-based regression across the iterations were averaged for each participant.

### Multidimensional scaling analysis

To visualize the representational geometry of WM, we performed multidimensional scaling (MDS). For each subject, ROI, TR, and location, we trained and tested a classifier to decode each of the 10 spatial locations. Training was performed in two different manners. First, we trained and tested on a TR-by-TR basis as we did with the cross-hemifield decoding analyses. This allows examination of the dominant geometry over time. Second, we trained on the last two time points of the post-cue delay (12 and 14 seconds) as we did with the temporal generalization decoding analyses. This allows emphasis on mnemonic codes. In each case, for each of the 10 spatial locations, classifier evidence was obtained for the 10 spatial locations and averaged across the time points of the pre-cue and post-cue delays, respectively. This resulted in a 10 x 10 matrix indicating the confusability of each spatial location for each delay period. All participant-level pre- and post-cue matrices were then concatenated (150 X 10 pre- and post-cue matrices), respectively, and used to calculate 10 X 10 correlation distance matrices. These correlation distance matrices were submitted to MDS using the cmdscale function implemented in R. The first three dimensions accounting for 70.7-96.5% variance (**Figures S5a and S5d**) were visualized. Each location was projected onto a 3-dimensional space showing the WM representational spaces of the ROIs by time (**Figures 6b, and 6c**). To statistically quantify the WM representational geometry, we repeated the same procedures separately at the individual participant level. Then, for each participant, we calculated the Euclidean distances between 10 spatial locations based on the 3-dimensional coordinates of spatial locations, resulting in a Euclidean distance matrix (Top row in **Figures S5g and S5h**). Since we supposed that the WM representational geometry might reflect either retinotopic or sequence position space, we estimated a matrix representing each space, using the visual angle distance of spatial locations (Bottom row in **Figures S5g and S5h**). In order to measure the similarity between the Euclidean distance and visual angle distance matrices, we calculated the Pearson’s correlation between the vectorized upper triangle portion of the Euclidean distance and visual angle distance matrices (excluding the diagonal) and performed Fisher’s Z transformation for the statistical quantification.

### Statistical analysis

Statistical tests for decoding analyses were based on permutation testing with 1000 iterations. Permutation testing avoids issues that can occur when testing against theoretical chance distributions at small sample sizes (Combrisson and Jerbi, 2015). For each iteration, the classifier was trained and tested with shuffled decoding labels, resulting in 15 null decoding accuracies (1 per participant) for each time point (0, 2, 4, 6, 8, 10, 12, and 14 sec) and each ROI (left/right IPS and VC). These null decoding accuracies were submitted to a one-tailed one-sample t-test against chance (one-tailed since decoding accuracy is not expected to be significantly below chance) and two-way repeated-measures ANOVAs, resulting in null T- or null F-values, which formed null distributions (**Figures 2 and 3**). Note that for t-tests, this procedure is identical to comparing actual decoding accuracy to a null distribution of decoding accuracies, but generalizes more easily to repeated-measures ANOVAs. P-values were the percentage of values in the null distribution that were greater than or equal to the observed T- or F-values obtained from the intact data, resulting in 0.001 as the lower limit for the P-value. We applied this calculation to other permutation statistical tests. (Note that for two-tailed tests, P-values were calculated based on the null distribution of the absolute null T-value and the absolute observed value). Similar to the above, statistical tests on beta weights obtained from RSA shown in Figure 5 were based on permutation testing over 1000 iterations. For each iteration, the shuffled correlation coefficient RSA matrix was submitted to model-based regression:

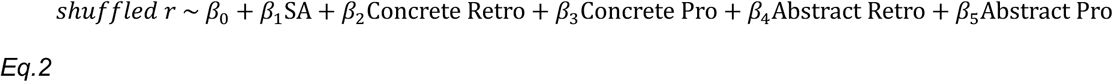

**Figure 2.**
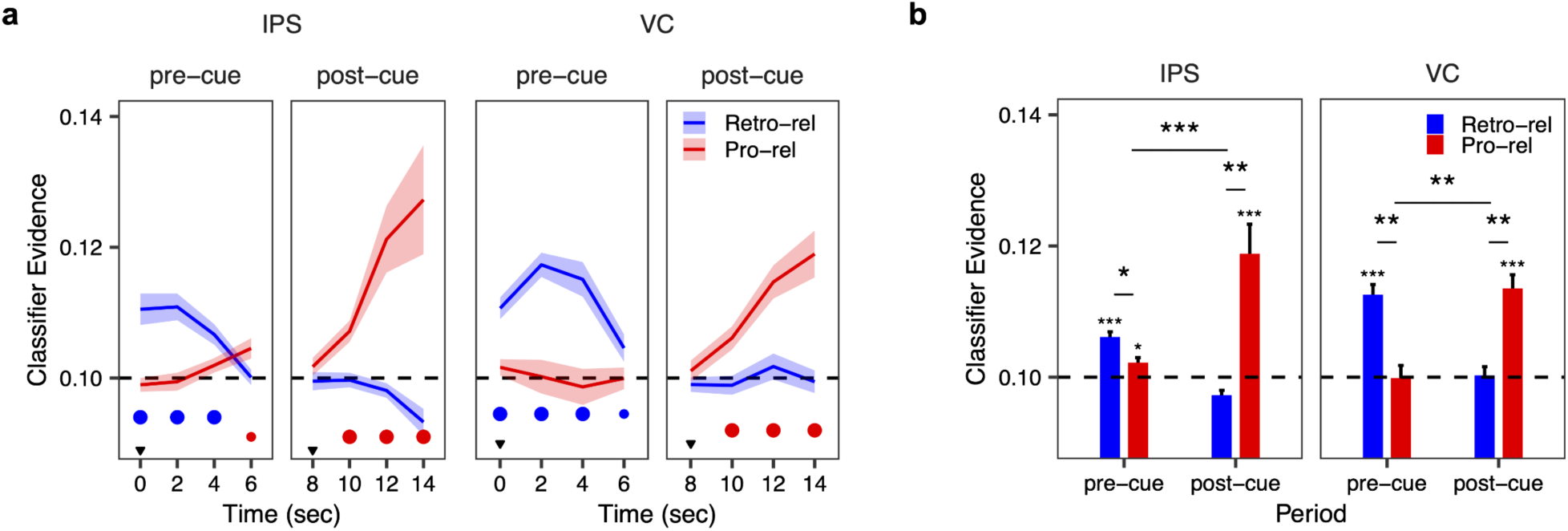
Tracking retrospective and prospective codes over time. **(a)** Classifiers were trained using data from the end of the post-cue delay (12 and 14 seconds) and then tested at each time point. Retrospective-relevant (‘Retro-rel’) refers to the classifier evidence corresponding to the spatial location of the just-probed item. Prospective-relevant (‘Pro-rel’) refers to the classifier evidence corresponding to the next spatial location following the reference (location following the previous probe in the pre-cue delay; location following the previous probe of the cued hemifield in the post-cue delay). The significance at each time point is denoted by small and medium dots, representing q < 0.05 and q < 0.01, respectively (FDR corrected). The inverted triangles denote the time points of the presentation of the probe and the cue, respectively. **(b)** Average of classifier evidence in each delay period (from 2 sec to 6 sec, ‘pre-cue’ period; from 10 sec to 14 sec, ‘post-cue’ period) for each ROI. *p < 0.05, **p < 0.01, ***p < 0.001. Error bars represent ±1 standard error of mean.

**Figure 3.**
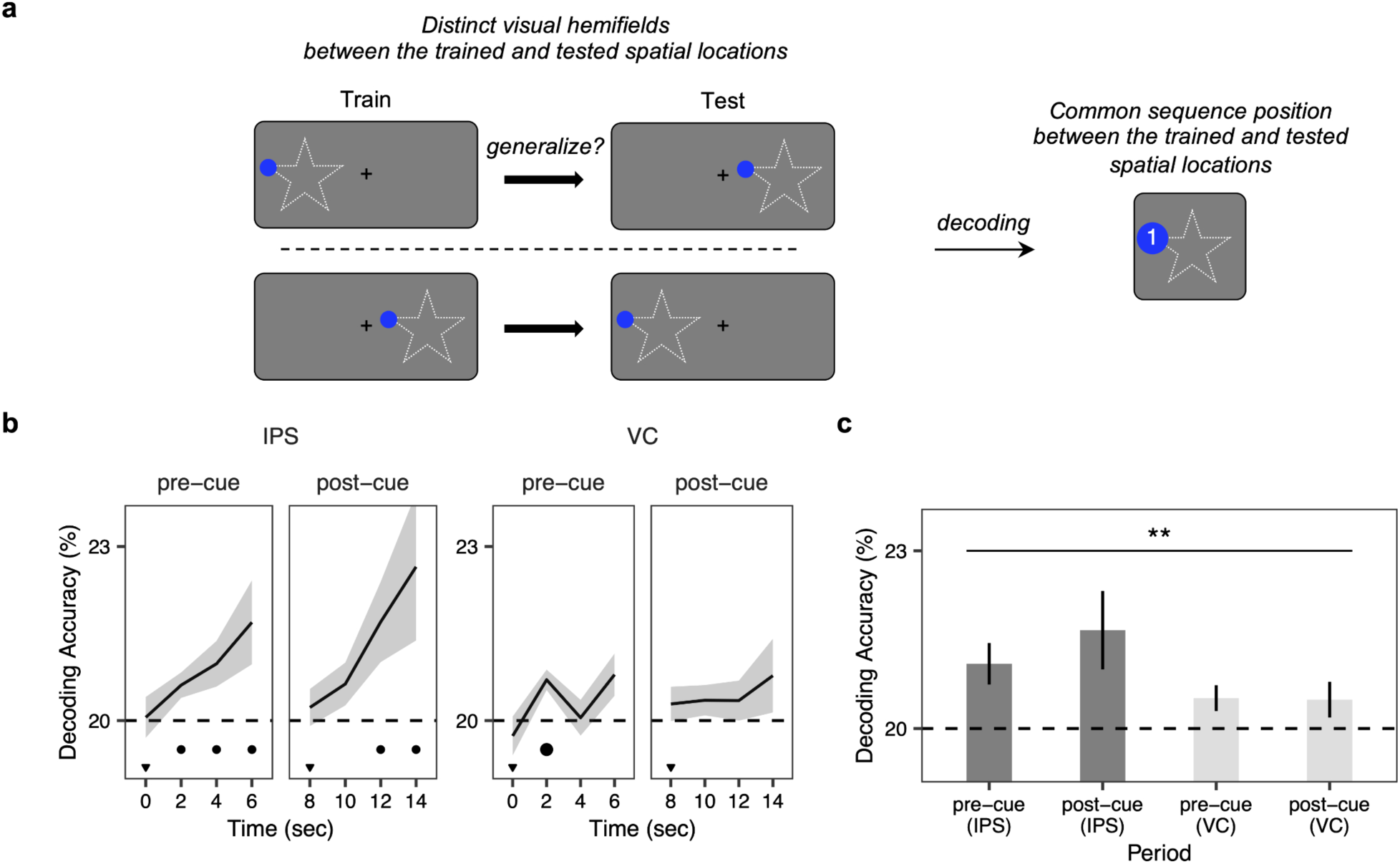
Decoding abstract sequence positions. **(a)** Schematic illustration of the cross-visual hemifield decoding analysis. Abstract coding was operationally defined based on generalization of decoding across visual hemifields. **(b)** The result of the cross-visual hemifield decoding analysis, in which training and testing the classifier were performed on different visual hemifields, respectively. The significance at each time point is denoted by small, and medium dots, representing q < 0.05, and q < 0.01, respectively (FDR corrected). The shaded regions represent ±1 standard error of mean. The inverted triangles in black denote the time points of the presentation of the probe stimulus and the cue, respectively. **(c)** Average of decoding accuracies from ‘pre-cue’ (2 sec to 6 sec) and ‘post-cue’ (10 sec to 14 sec) period in each ROI. The horizontal dashed line indicates the chance level of 20%. **p < 0.01. Error bars represent ±1 standard error of mean.

For each iteration, we obtained 15 null beta weights (1 per subject) for each time point, each ROI, and each regressor (Concrete Retro, Concrete Pro, Abstract Retro, and Abstract Pro). These null beta weights were submitted to one-tailed one-sample t-tests against a value of 0 (one-tailed because only beta weights above a value of 0 are expected), two-tailed paired t-tests and a two-way RM ANOVA, resulting in null T- or null F-values, which formed the null distributions (**Figures 5a and 5b**).

The statistical test on the brain-behavior correlation analysis was based on a permutation test with 1000 iterations. For each iteration, we shuffled the participant order of 15 observed averaged beta weights of interest and correlated it with the intact participant order of behavioral data, resulting in the null distribution of the null T-values (**Figure 5c**).

Statistical test on MDS shown in Figure 6 was based on permutation testing with 1000 iterations. Since the output of the cmdscale function in R was used as the coordinates corresponding to WM representations for 10 spatial locations in 3D space, we shuffled the outputs but kept the label of 10 spatial locations intact for each iteration. Using the shuffled multidimensional 3D coordinates, we calculated the null Euclidean distances between 10 spatial locations, generating a null Euclidean distance matrix. These matrices were then correlated with either the intact retinotopic matrices or sequence position matrices by vectorizing the upper triangular portion of each matrix. For the statistical test, the correlation value was Fisher’s Z transformed, resulting in 15 null Fisher’s Z values (1 per subject) for each time point, each ROI, and each space (retinotopic and position). These null Fisher’s Z values were submitted to two-tailed paired t-tests, and two-way/three-way RM ANOVAs, resulting in the null distribution of the null T- and F-values (**Figures 6d and 6e**).

For all statistical tests across multiple time points, multiple comparisons were corrected using false-discovery rate (FDR) (Benjamini and Hochberg, 1995).

### Data and code availability

Preprocessed data and all original code have been deposited at Open Science Framework and are publicly available at https://osf.io/n5ake/ as of the date of publication.

## Results

Participants separately and continuously tracked spatial locations in the left and right visual hemifields (**Figure 1b**). On each trial, a cue indicated the relevant hemifield for the forthcoming probe affording a prospective prediction of the sequencing-matching location. Participants indicated whether the spatial location of the probe followed the spatial location of the last stimulus in the cued hemifield (reference). Half of the probes were matches while half were non-matches with non-match probes equally drawn from the three non-matching spatial locations excluding the reference (i.e., there were no stimulus repeats).

After responding, the spatial location of the probe became the reference for the next probe of the same hemifield. The cue was separated from the previous (pre-cue delay) and forthcoming (post-cue delay) probes allowing for periods to track WM codes of different forms (retrospective vs. prospective, concrete/location vs. abstract/position; **Figure 1c**).

Behavioral data indicated that participants performed the task well (mean accuracy = 94.8 %). To verify that participants appropriately prioritized relevant information (i.e., the cued hemifield), we separately characterized trials in which the cue indicated the same visual hemifield as the previous trial (‘Repeat’) or the opposite visual hemifield (‘Switch’). As anticipated, reaction times (RT) and error rates (ER) were larger in ‘Switch’ (610.1 ms, 7.4 %) relative to ‘Repeat’ trials (594.5 ms, 3.1 %; *t*_(14)_ = 2.53, *p* = .024 for reaction time; *t*_(14)_ = 3.44, *p* = .004 for error rate, paired *t*-tests; **Figure 1d**) consistent with a cost for switching priority. These data confirm that participants performed the task as expected.

### Location codes are transformed over time

We hypothesized that WM codes would transform over time trading off from retrospective to prospective (**Figure 1c**). Some work suggests that sensory and mnemonic signals are multiplexed in the visual cortex and IPS (Rademaker et al., 2019; Chunharas et al., 2024). In order to focus on mnemonic representations, classifiers were trained on patterns drawn from the end of the post-cue delay (12 and 14 seconds post-stimulus onset) and then tested across time points from 0 to 14 seconds (see **Figure S4a** for classifiers trained at the end of the pre-cue delay). Classifiers were trained and tested separately for each region-of-interest (ROI; left and right IPS and VC) to discriminate the activation patterns corresponding to each of the 10 spatial locations.

If WM codes transform over time, the location held in mind at the end of the post-cue delay just prior to the probe (i.e., prospective) would differ from the location encoded during and shortly after stimulus presentation (i.e., retrospective). Therefore, we labeled each training pattern as though it coded for the cued, prospective location. Then we tested the classifier separately for each time point. We separately detailed classifier evidence for the retrospective spatial location in the relevant hemifield (retrospective-relevant; ‘Retro-rel’), the retrospective spatial location in the irrelevant hemifield (retrospective-irrelevant; ‘Retro-irrel’), the prospective spatial location in the relevant hemifield (prospective-relevant; ‘Pro-rel’), and the prospective spatial location in the irrelevant hemifield (prospective-irrelevant; ‘Pro-irrel’; **Figure 1c**). Relevance refers to the just-probed hemifield in the pre-cue period and the cued hemifield in the post-cue period. The results were averaged across hemispheres for each of the IPS and VC, respectively.

**Figure 2** depicts the classifier evidence for the retrospective-relevant (‘Retro-rel’) and prospective-relevant (‘Pro-rel’) locations. In both the IPS and VC, classifier evidence for the retrospective-relevant spatial location was significantly above chance levels during the pre-cue delay, whereas it dropped to chance during the post-cue delay (note, all statistical tests are performed against a permutation null (Combrisson and Jerbi, 2015); see Methods). By contrast, classifier evidence for the prospective-relevant spatial location showed the opposite pattern (Figure 2a). Separating the data by post sequence-match (**Figure S1a**) and post sequence-non-match (**Figure S1b**), and by repeat (**Figure S1c**) and switch (**Figure S1d**) trials revealed that these patterns were shaped, but not driven by carry-over from prior trials (sequence-match/non-match) or periods (repeat/switch). To further quantify these patterns, we averaged each classifier evidence within each delay period (from 2 sec to 6 sec, ‘pre-cue’; from 10 sec to 14 sec, ‘post-cue’) and submitted the data to a 2 (period: ‘pre-cue’ vs. ‘post-cue’) x 2 (location: ‘Retro-rel’ vs. ‘Pro-rel’) repeated-measures ANOVA for each of the IPS and VC (Figure 2b). We observed a significant interaction of period and location in both the IPS (*F*_(1, 14)_ = 19.35, *p* < .001) and VC (*F*_(1, 14)_ = 16.33, *p* < .01) confirming a shift from retrospective to prospective coding in each area.

To examine whether the timing of these transitions differed by region, we performed ROI X location repeated-measures ANOVA at individual time points. Significant interactions were observed at 2, 4, and 6 seconds (*F*_(1, 14)_ > 4.9, *p*s < .01). Follow-up paired t-tests revealed that the VC showed stronger evidence for the retrospective relevant item than the IPS at 2 and 4 seconds (*ts*_(14)_ > 3.31, *p*s < .01), whereas the IPS showed stronger evidence for the prospective relevant item than the VC at 6 seconds (*t*_(14)_ = 2.66, *p* = .019). These data suggest that although both areas transition from coding retrospective to prospective items, the VC more strongly codes for retrospective locations during the early delay, while the IPS shifts towards coding for prospective locations earlier than the VC.

Most evidence for either the retrospective-irrelevant or prospective-irrelevant spatial locations were at or under chance (**Figure S2**), indicating that decodable representations in the IPS and VC were restricted to prioritized items as consistent with substantial past work (Lewis-Peacock et al., 2012; Sprague et al., 2016; LaRocque et al., 2017; Lorenc et al., 2020; Yu et al., 2020). In subsequent analyses, we will therefore focus on relevant items.

Taken together, the results indicate that WM was transformed over the delay periods from retrospectively representing the past to prospectively representing expectations for the future.

### Abstract sequence information is coded in the IPS

In the current task, anticipation of prospective spatial locations in each visual hemifield was guided by the same abstract spatial sequence. Abstraction is commonly defined as generalization across contexts (Bernardi et al., 2020; Kwak and Curtis, 2022; Duan and Curtis, 2024). Here, each visual hemifield formed a distinct context. To test for abstract coding of sequence, we examined the extent to which representations generalized across hemifields by training a classifier using representations from one hemifield and testing it with representations from the other hemifield (cross-visual hemifield decoding; Figure 3a). Training and testing were performed separately for each TR using labels of the relevant sequence position. We found significant cross-visual hemifield decoding during both the pre-cue and post-cue delays in the IPS with decoding accuracy ramping up over time during both phases (Figure 3b). By contrast, in the VC, the classifier showed above-chance decoding accuracy only during a single time point in the pre-cue delay. To better quantify this result, we averaged the decoding accuracies from 2 sec to 6 sec to form the ‘pre-cue’ period and from 10 sec to 14 sec to form the ‘post-cue’ period separately for each region, and submitted the data to a region (IPS, VC) x period (‘pre-cue’, ‘post-cue’) repeated-measures ANOVA. Abstract sequence position was more strongly decoded in the IPS relative to the VC (main effect of region *F*_(1, 14)_ = 8.74, *p* = .008; Figure 3c). Neither the main effect of period nor the interaction between period and region was significant. Collectively, although there was some weak evidence for abstract codes in the VC, abstract sequence information was prominent across delay periods in the IPS.

### Concrete and abstract codes of working memory are transformed over time

To better examine the extent to retrospective, prospective, concrete, and abstract codes are multiplexed versus trade off over time, we turned to representational similarity analysis (RSA) (Kriegeskorte et al., 2008). For each of the 10 spatial locations, we extracted the patterns of activation in each of our 4 ROIs (left and right IPS and VC). Training patterns were formed by averaging over the last two time points of the post-cue delay as we did above (see **Figure S4b** for training during the pre-cue delay). Then in a testing set, for each time point, we calculated the correlation similarity between the activity pattern at that time point and each training pattern (Figure 4a). This was done separately for each ROI. We then decomposed representations in the IPS and VC into 1) coarse representations of the attended hemifield (spatial attention), 2) fine-grained representations of the retrospective, relevant spatial location (concrete retrospective), 3) fine-grained representations of the prospective, relevant spatial location (concrete prospective), 4) abstract representations of the retrospective sequence position (abstract retrospective), and 5) abstract representations of the prospective sequence position (abstract prospective) using multiple linear regression. Each regressor represents a representational similarity matrix corresponding to a hypothetical pure representation (see Methods; Figure 4a). The regressors incorporated distance-dependent similarity based upon normalized visual angle (i.e., nearby locations are more similar to one another than far locations). Concrete regressors were formed by training/testing in the same hemifield, while abstract regressors were formed by training/testing across hemifields. Based on the analyses above, we expected that retrospective codes would transition to prospective codes, both concretely and abstractly (Figure 4b).

**Figure 4.**
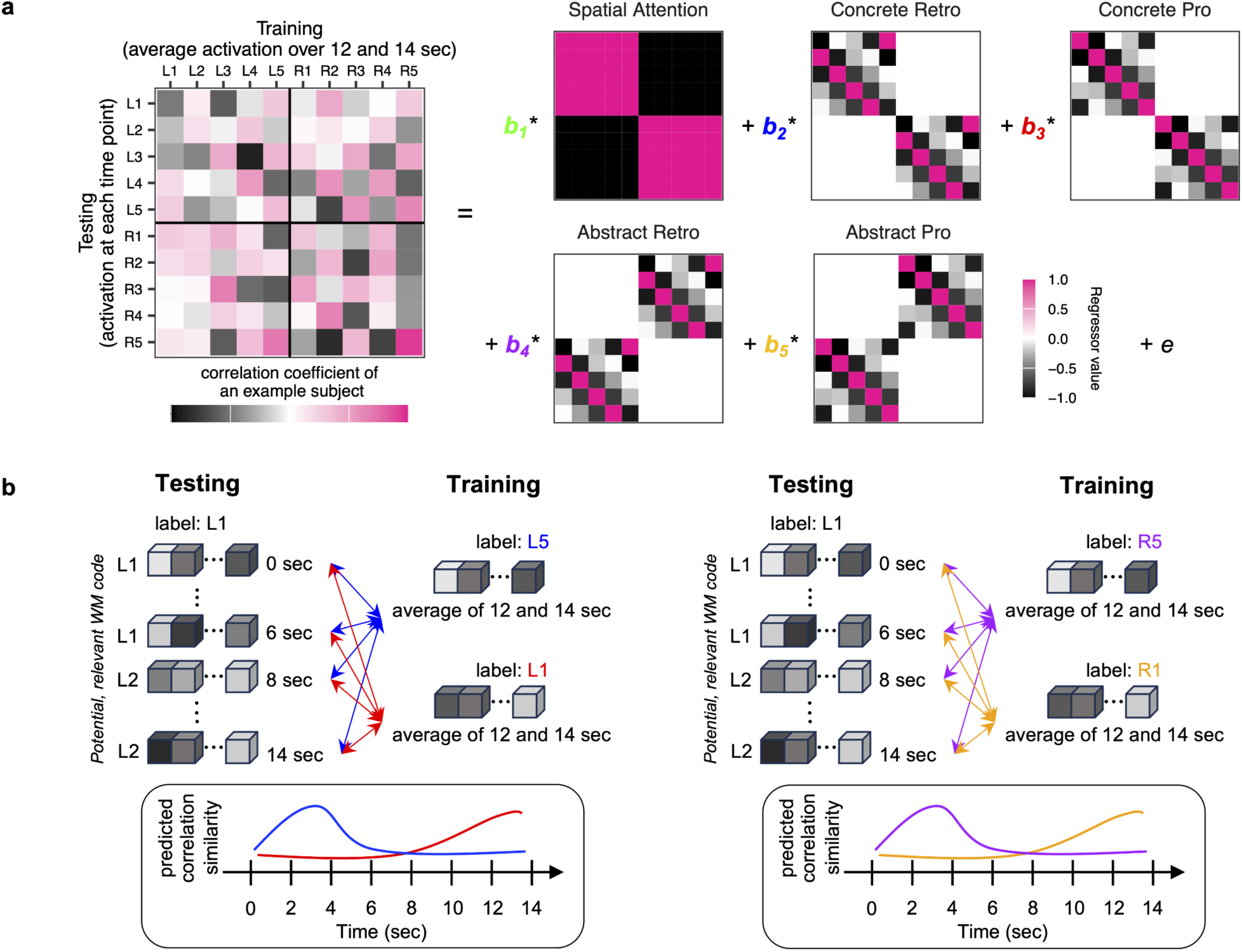
Schematic illustration of the representational similarity analysis. **(a)** The left depicts an example representational similarity matrix formed by correlating training patterns of each location with testing patterns of each location. These matrices were submitted to multiple linear regression using regressors denoting idealized matrices of spatial attention, concrete retrospective coding, concrete prospective coding, abstract retrospective coding, and abstract prospective coding. We estimated the beta weight for each regressor at each time point. Training patterns were formed by averaging the activation patterns at 12 and 14 seconds post-stimulus onset. Hypothetical models were created assuming that the training patterns reflect the prospective location/position. Hence, retrospective models reflect a correspondence between testing patterns (e.g., L1, the first row) that correspond to the location/position preceding the training pattern (e.g., L5) with increasing dissimilarity as a function of increasing retinotopic distance. **(b)** Schematic of the predicted correlation similarities representing a transition between concrete retrospective to concrete prospective coding (left) and abstract retrospective to abstract prospective coding (right). For example, the training pattern of L5 (or R5 if abstract coding) was assumed to have the highest similarity with the testing pattern of L1 during the ‘pre-cue’ delay period. By contrast, the training pattern of L1 (or R1 if abstract coding) was assumed to have the highest similarity with the testing pattern of L1 during the ‘post-cue’ delay period.

**Figure 5.**
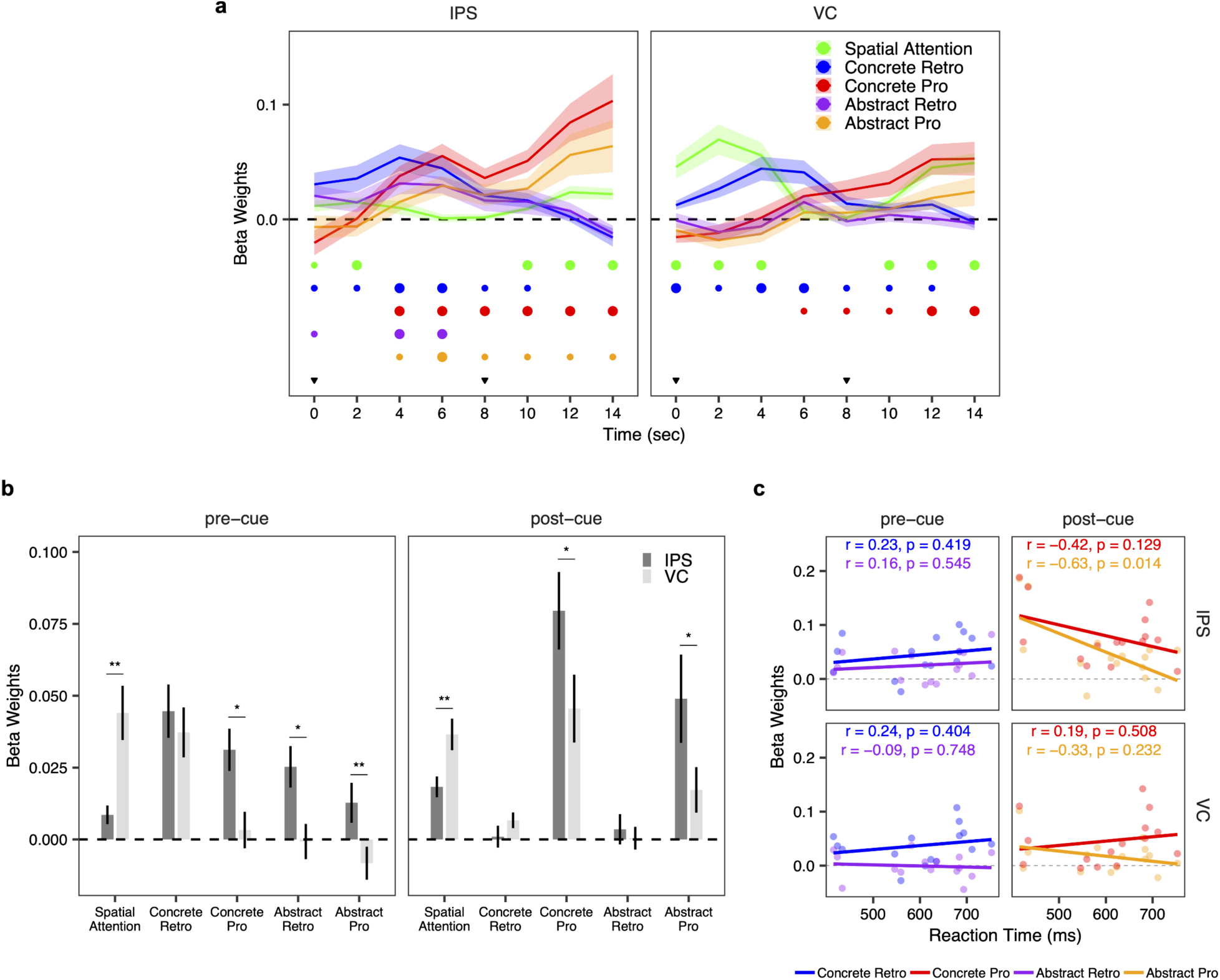
Simultaneous and dynamic coding of the past and future. **(a)** The estimated beta weights for each regressor at each time point. The beta weights were compared against a value of 0 and corrected for multiple comparisons using false discovery rate (FDR). The significance at each time point is denoted by small and medium dots, representing q < 0.05 and q < 0.01, respectively (FDR corrected). The shaded regions represent ±1 standard error of mean. The inverted triangles in black denote the time points of the presentation of the probe stimulus and the cue, respectively. **(b)** To contrast the IPS and VC, spatial attention and each WM code were separately formed by averaging beta weights during the ‘pre-cue’ period (2 sec to 6 sec) and the ‘post-cue’ period (10 sec to 14 sec) in each ROI. Error bars represent ±1 standard error of mean. *p < 0.05, **p < 0.01. **(c)** Correlations between the strengths of WM codes and RT for each ROI. Each dot represents each participant.

**Figure 6.**
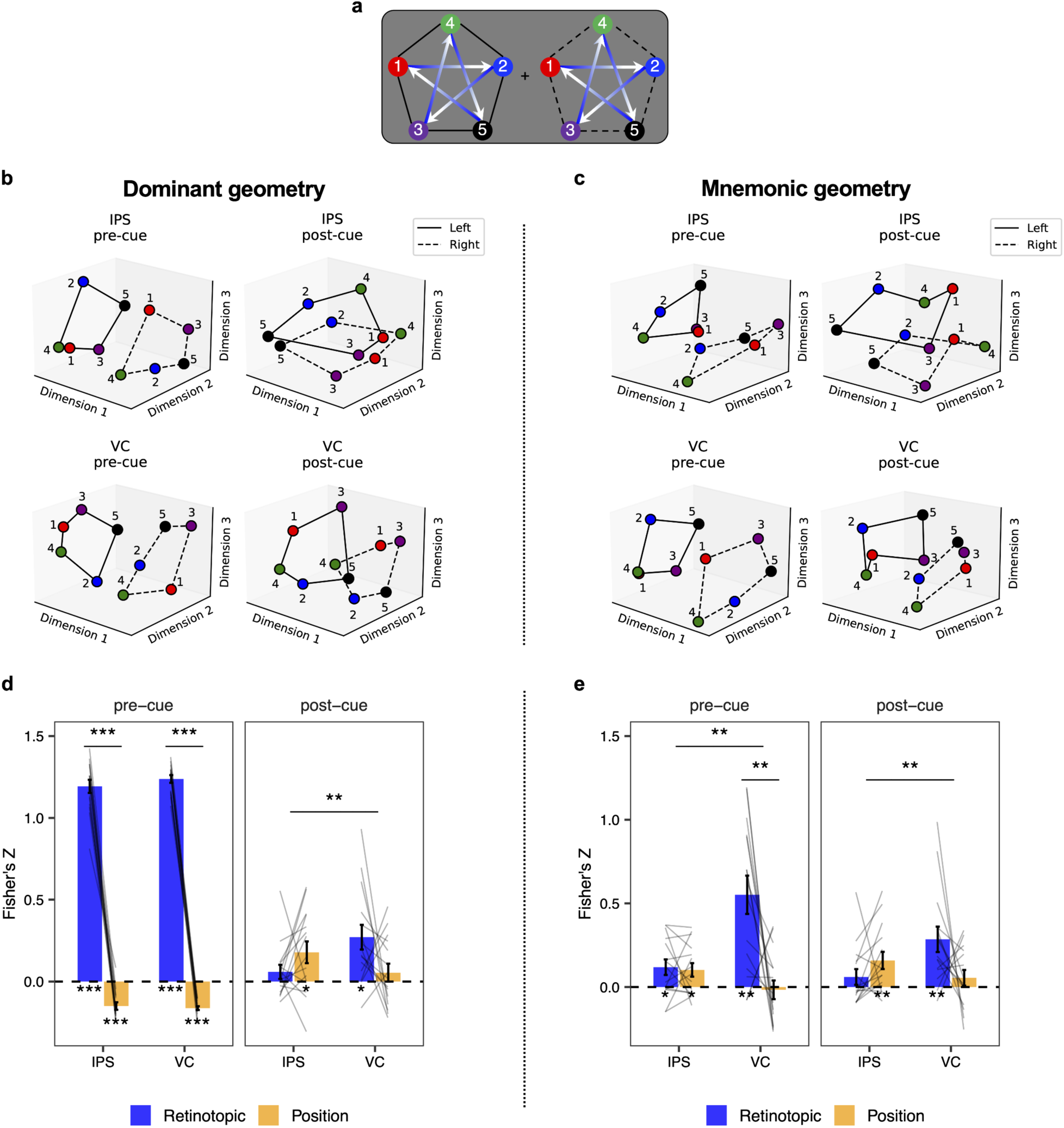
Representational geometry of WM. **(a)** Schematic illustration of the spatial locations. **(b)** Each of the 10 spatial locations is plotted by the first three dimensions of the group-level dominant MDS geometry. Spatial locations are separately color coded by sequence position. Spatial locations in the left and right visual hemifields are connected by solid and dotted black lines, respectively. **(c)** Each of the 10 spatial locations is plotted by the first three dimensions of the group-level mnemonic MDS geometry. Spatial locations are separately color coded by sequence position. Spatial locations in the left and right visual hemifields are connected by solid and dotted black lines, respectively. **(d)** The result of the correlation between Euclidean distance among locations calculated in retinotopic space (blue) and position space (orange) and the dominant MDS geometry space. Note that the correlation analysis was performed at the individual participant level. Each line on the bar graph represents each participant. *p < 0.05, **p < 0.01, ***p = 0.001. Error bars represent ±1 standard error of mean. **(e)** The result of the correlation between Euclidean distance among locations calculated in retinotopic space (blue) and position space (orange) and the mnemonic MDS geometry space. Note that the correlation analysis was performed at the individual participant level. Each line on the bar graph represents each participant. *p < 0.05, **p < 0.01, ***p = 0.001. Error bars represent ±1 standard error of mean.

As shown in Figure 5a, we found evidence consistent with these predictions. In both the IPS and VC, a concrete retrospective code emerged early during the pre-cue delay, but diminished over time until it was absent by the end of the post-cue delay. The fall of concrete retrospective codes was mirrored by rising concrete prospective codes which started before the cue (van Ede et al., 2021) and peaked at the end of the post-cue delay. In the IPS, we observed a similar tradeoff in abstract codes from retrospective to prospective. However, abstract codes were not significantly observed in the VC. Interestingly, abstract and prospective codes emerged at the same time point in the IPS (4 second post-stimulus onset) and preceded the onset of concrete prospective codes in the VC (6 seconds post-stimulus onset). Although the sluggishness of the BOLD signal precludes detailed timing information, the data are consistent with the use of abstract sequence position to inform concrete expectations of the future. In other words, the stimulus spatial location (concrete retrospective) may be translated into an abstract sequence position (abstract retrospective) which can then be transformed into the next abstract sequence position (abstract prospective), affording a prediction of the probe spatial location (concrete prospective).

To further quantify these observations, for each region, we averaged the beta weights from 2 sec to 6 sec to form the ‘pre-cue’ period, and directly contrasted the IPS and VC with paired t-tests (Figure 5b). Spatial attention was more strongly represented in the VC than IPS (*t*_(14)_ = -3.85, *p* = .006), whereas concrete retrospective coding did not differ between the regions (*t*_(14)_ = 0.77, *p* = .46). By contrast, the IPS showed stronger coding of the abstract retrospective (*t*_(14)_ = 2.79, *p* = .013), concrete prospective (*t_(14)_* = 3.05, *p* = .014), and abstract prospective (*t*_(14)_ = 2.81, *p* = .008) codes in the pre-cue period. These data indicate that abstract and prospective codes emerge earlier in the IPS than VC. Next, for each region, we averaged the beta weights from 10 sec to 14 sec to form the ‘post-cue’ period. These data indicated that spatial attention remained stronger in the VC than IPS (*t*_(14)_ = -3.88, *p* = .003), whereas concrete (*t*_(14)_ = 2.84, *p* = .013) and abstract (*t*_(14)_ = 2.59, *p* = .025) prospective coding remained stronger in the IPS than the VC (Figure 5b). Finally, we cross-correlated the magnitudes of prospective codes during the post-cue period. Abstract prospective codes in the IPS were correlated with concrete prospective codes in the IPS (*r* = 0.75, *p* = 0.005), which were, in turn, correlated with concrete prospective codes in the VC (*r* = 0.56, *p* = 0.025). Hence, these data are consistent with a transformation and relay of codes supporting WM.

Next, we examined how representational codes were related to behavioral performance. We reasoned that behavioral preparedness (i.e., faster RTs) would be related to prospective coding, rather than retrospective coding. Consistent with this idea, RT was negatively related to abstract prospective coding during the post-cue period in the IPS (*r* = -0.63, *p* = 0.014), but not VC (*r* = -0.33, *p* = 0.232). A similar, non-significant trend was observed for concrete prospective coding in the IPS (*r* = -0.42, *p* = 0.129). However, this trend appeared to be driven by shared variance with abstract coding (described above) as partialling out concrete prospective coding left the relationship between abstract prospective coding in the IPS and RT largely intact (r = -0.5264, p = 0.055) while the reverse was not true (r = 0.1029, p = 0.72). Moreover, concrete prospective coding in the VC was unrelated to RT and reversed in sign (*r* = 0.19, *p* = 0.508). Similarly, concrete and abstract retrospective coding during the pre-cue period was non-significantly related to RT (*ps* > 0.3). These results were replicated when using inverse efficiency indicating that the results could not be attributed to a speed-accuracy tradeoff (**Figure S3**). Collectively, these data suggest that abstract prospective coding in the IPS facilitates future behavioral performance.

### WM representational space is reorganized over time

The RSA analyses revealed multiplexing of representational codes supporting WM in both the IPS and VC. To more directly visualize these representations, we turned to multidimensional scaling (MDS) which helps to elucidate representational geometry. For each delay period, region, and time point, we submitted the similarity structure of the 10 spatial locations to MDS and plotted each location in the space formed by the first three dimensions (see Methods). We performed this analysis in two ways. First, we trained and tested on a TR-by-TR basis as we did with the cross-hemifield decoding analysis. This allows visualization of the dominant geometry over time (dominant geometry). Second, we trained on the end of the post-cue delay (12 and 14 second post-stimulus onset) as we did with the temporal generalization and representational similarity analyses. This allows honing in on mnemonic codes (mnemonic geometry).

Each analysis revealed a clear geometry during each delay period, and transformation of the representations supporting WM over time. Starting with the dominant geometry, during the ‘pre-cue’ period, representations were separated by hemifield along the first dimension in both the IPS and VC (Figure 6b **and Figure S5b**). For both areas, a clear circular geometry was present across the second and third dimensions resembling the structure of the visually-presented locations (Figure 6b **and Figure S5c**). Across hemifields these circular structures were vertically aligned but mirror-reversed consistent with distance in retinotopic space (e.g., position 2 in the left hemifield is close to position 1 in the right hemifield; Figure 6a). Hence, retinotopic space was well-recapitulated across the three dimensions of MDS in both the IPS and VC. Similar results were obtained in the VC for the mnemonic geometry (Figure 6c). In the IPS, the mnemonic geometry was also split by hemifield. However, in this case, the sequence positions across the hemifields were aligned along the second and third dimensions. Thus, the mnemonic geometry revealed evidence of abstraction. These data indicate that retinotopic and abstract sequence codes were multiplexed in the IPS, while the VC represented retinotopic codes.

In the ‘post-cue’ period, the dominant geometry and mnemonic geometry were similar to one another in the IPS and VC, respectively, but differed across regions. In the IPS, hemifield discrimination was reduced and relegated to the third dimension (Figures 6b **and 6c**). A circular geometry resembling the star-sequence was observed along the first and second dimensions (**Figures S5b and S5e**). These circular geometries were aligned by position across hemifields. In VC, hemifield discrimination remained prominent along the first dimension. Alignment by sequence position was also observed in VC, albeit across the second and third dimensions (**Figures S5c and S5f**). That is, the ‘post-cue’ geometry in the VC resembled the ‘pre-cue’ mnemonic geometry of the IPS. Hence, representational transformations were observed in both the IPS and VC transitioning away from a concrete, retinotopic codes towards abstract, position codes. However, the areas differed in their principal coding axis. The IPS more prominently coded for sequence position while the VC more prominently coded for hemifield.

To quantitatively evaluate the representational dynamics in the multidimensional space, we examined the extent to which Euclidean distance among representations in each of the dominant and mnemonic geometries resembled retinotopic distance and abstract position distance. Retinotopic distance was calculated as the distance in visual angle among spatial locations. Position distance was calculated similarly, but assumed that positions across hemifields were identical (**Figures S5g and S5h**). For each subject, we correlated the matrix of Euclidean distances in each MDS geometry with the matrix of visual angle distances in retinotopic and position space, respectively, separately for the pre- and post-cue delays. Correlation values were z-transformed and submitted to a period (‘pre-cue’ vs. ‘post-cue’) by region (IPS vs. VC) by space (retinotopic vs. position) repeated-measures ANOVA (Figures 6d **and 6e**).

For the dominant geometry, we found a significant three-way interaction (*F*_(1, 14)_ = 5.44, *p* = .035; see Table 1 for complete stats). Specifically, in the ‘pre-cue’ period, the representational geometry closely resembled retinotopic space much more so than position space in both the IPS and VC (region X space repeated-measures ANOVA in the ‘pre-cue’ period; main effect of space; *F*_(1, 14)_ = 2181.08, *p* = .001). By contrast, in the ‘post-cue’ period, the IPS representational geometry was more similar to the sequence position space than the retinotopic space, whereas the reverse was true in the VC representational geometry (region X space repeated-measures ANOVA in the ‘post-cue’ period; region x space interaction: *F*_(1, 14)_ = 12.72, *p* = .004). For the mnemonic geometry, a three-way interaction was not significant (*F*_(1, 14)_ > 1.63, *p* = .222; see Table 2 for complete stats). Instead, a significant region X space interaction was observed (*F*_(1, 14)_ = 14.05, *p* = .003) driven by more prominent retinotopic than abstract position coding in the VC in both the ‘pre-cue’ and ‘post-cue’ delays with the opposite true of the IPS.

**Table 1.** Repeated-measures ANOVA results associated with Figure 6d.

| RM ANOVA | Factors | *F* | *p* |
| --- | --- | --- | --- |
| 3-Way | Space | 180.750 | 0.001 |
|  | Period | 116.943 | 0.001 |
|  | Region | 2.233 | 0.151 |
|  | Space X Period | 297.681 | 0.001 |
|  | Region X Space | 14.571 | 0.003 |
|  | Period X Region | 0.610 | 0.465 |
|  | Space X Period X Region | 5.439 | 0.035 |
| 2-Way ('pre-cue' period) | Space | 2181.076 | 0.001 |
|  | Region | 0.720 | 0.429 |
|  | Region X Space | 0.943 | 0.335 |
| 2-Way ('post-cue' period) | Space | 0.312 | 0.579 |
|  | Region | 1.781 | 0.218 |
|  | Region X Space | 12.716 | 0.004 |

Fisher transformed correlation value was submitted to each RM ANOVA.

**Table 2.** Repeated-measures ANOVA results associated with Figure 6e.

| RM ANOVA | Factors | *F* | *p* |
| --- | --- | --- | --- |
| 3-Way | Space | 7.902 | 0.014 |
|  | Period | 3.318 | 0.083 |
|  | Region | 21.762 | 0.001 |
|  | Space X Period | 5.783 | 0.029 |
|  | Region X Space | 14.045 | 0.003 |
|  | Period X Region | 2.864 | 0.104 |
|  | Space X Period X Region | 1.631 | 0.222 |
| 2-Way ('pre-cue' period) | Space | 14.498 | 0.001 |
|  | Region | 10.659 | 0.004 |
|  | Region X Space | 9.113 | 0.007 |
| 2-Way ('post-cue' period) | Space | 0.651 | 0.433 |
|  | Region | 9.431 | 0.013 |
|  | Region X Space | 11.493 | 0.003 |

Fisher transformed correlation value was submitted to each RM ANOVA.

Taken together, these results suggest that representational geometries in the IPS and VC were organized in a different manner. Specifically, concrete, retinotopic and abstract, sequence position codes were multiplexed in the ‘pre-cue’ delay in the IPS. Over time, retinotopic coding gave way to abstract sequence position codes. By contrast, VC representational geometry was dominated by a concrete, retinotopic code across delays.

## Discussion

Although the primary function of WM is to guide future cognition (Nobre and Stokes, 2019), the mechanisms supporting the transformation of past sensory codes into abstract codes that prepare for the future have been elusive. Here, we found evidence for concrete, retrospective codes in the VC and the IPS when human participants maintained past sensory information (pre-cue delay). Over time concrete, retrospective codes disappeared concomitant with the appearance of prospective codes. This transition was marked by the emergence of abstract sequence position codes in the IPS. Moreover, the strength of abstract prospective coding in the IPS was related to more efficient behavioral performance consistent with a signal that prepares for future cognition. Lastly, dimensionality reduction (MDS) revealed how information is multiplexed and transformed over time. Specifically, concrete, sensory and abstract sequence position codes were simultaneously present in the IPS early in the delay. Over time, sensory codes dissipated and abstract sequence position codes dominated. By contrast, representations in the VC remained more sensory-like both early and late. Taken together, these results detail how WM transforms to support future cognition with dissociable roles of VC more strongly representing concrete, sensory-like codes and the IPS more strongly representing the abstract future.

Numerous previous studies have revealed that concrete retrospective sensory signals can be decoded in the IPS and VC during WM delay periods (Harrison and Tong, 2009; Serences et al., 2009; Ester et al., 2015; Bettencourt and Xu, 2016; Sprague et al., 2016; Rademaker et al., 2019). These findings have led to the conclusion that both regions serve as critical sites for the storage of visual WM. However, a central function of WM is the transformation of past information into future-relevant information beyond just accurately reproducing or recognizing past information. Recent studies using electroencephalography or eye-tracking have provided evidence showing that anticipated actions (van Ede et al., 2019; Boettcher et al., 2021) and spatial locations (Gunseli et al., 2024; Liu et al., 2024) are prospectively coded during WM delays. Furthermore, human fMRI studies have decoded prospective category-level WM representations in areas processing visual objects such as fusiform gyrus, middle temporal gyrus, and lateral occipital cortex (Lewis-Peacock and Postle, 2008; Lewis-Peacock et al., 2012). Building on these findings, our data show that the fine-grained representation of past spatial locations in the IPS and VC are transformed into anticipated spatial positions when preparing for a sequential probe. Moreover, by inspecting the geometry of these representations, we find evidence that representational codes become abstracted away from sensation such that locations that are retinotopically distant nevertheless align in representation space when they share meaningful features (Courellis et al., 2024). Changes in representational alignment over time may explain why training classifiers during WM results in superior decoding of WM relative to training classifiers on perception (Harrison and Tong, 2009; Rademaker et al., 2019; Lorenc et al., 2020). This result implies that the IPS and VC store not only the concrete past but also the anticipated future, with the future stored in a more abstract format than the past. These data are consistent with suggestions that abstraction facilitates planning for the future (Ho et al., 2019).

Although several works have shown similar contributions of the IPS and VC to visual WM, other work has suggested that their roles may be dissociable. For example, some studies have demonstrated that the presence of a distractor during a WM delay disrupts or biases WM representations in the VC, whereas WM representations in the IPS are robust against distractors (Bettencourt and Xu, 2016; Lorenc et al., 2018; Rademaker et al., 2019). This work implies that the IPS may store not only concrete, sensory-like WM codes but also non-stimulus-driven WM representations that are transformed away from the sensory signals (Rademaker et al., 2019). Consistent with this idea, our data show evidence of both concrete sensory and abstract position signals in the IPS with concrete representations transitioning into abstract representations over time. By contrast, evidence for such transitions in the VC were weaker or non-significant depending upon the exact form of the analysis. We suggest that the hints of such signaling in the VC may be due to top-down signals from the IPS. Such an idea is consistent with recent work showing that during WM delays, the IPS shapes the contents of VC through feedback (Xu, 2023), which could potentially be examined in future work by distinguishing representations across layers of VC (van Kerkoerle et al., 2017; Lawrence et al., 2018). Moreover, not only were abstract prospective codes consistently stronger in the IPS than VC, the strength of abstract prospective codes in the IPS were related to improved behavior. Collectively, these data indicate dissociable roles of the VC and IPS with the former involved in concrete, sensory representation and the latter important when abstract, prospective codes facilitate performance (Brincat et al., 2018). These data are broadly consistent with cortical progressions from sensory to more abstract coding (Mesulam, 1998; Brincat et al., 2018).

Recent work has shown that VC representations can also be abstracted away from a strict sensory format. VC representations supporting WM can generalize across gratings and motion directions into a line-like format (Kwak and Curtis, 2022), and can show invariance to aperture biases that affect sensory representations (Duan and Curtis, 2024). In those works, generalization was examined within the same spatial receptive field. Here, we examined generalization across spatial receptive fields (i.e., across visual hemifields). Hence, that abstract coding was weak to absent in the VC in our data does not undermine the presence of other forms of abstract coding in the VC. Rather, there may well be different levels of abstraction which transition across the visual hierarchy. Since we did not perform retinotopic mapping to carefully tease apart different levels of the visual hierarchy here examination of how abstract coding in the present task varies along the levels of the visual hierarchy is a question for future work.

According to a recent proposal, the nature of abstraction in WM is to transform its representational structure—reflecting past informational content—into a future-oriented format to guide the upcoming task (Myers et al., 2017; Wang et al., 2019; Xie et al., 2022; van Ede and Nobre, 2023). Panichello and Buschman (Panichello and Buschman, 2021) contrasted selection from WM with selection from perception (i.e., attention) in monkeys. They found that selection in both forms generalized in the prefrontal cortex (PFC), but not parietal cortex or VC, providing evidence for an abstract mechanism of control in the PFC. Moreover, PFC representations of WM items transformed from a pre-cue subspace wherein WM representations for different colors were multiplexed with location, to a post-cue subspace wherein representations for different colors aligned regardless of location. The same transformation in representational geometry was also observed in a recurrent neural network trained to carry out an equivalent cued-recall paradigm (Piwek et al., 2023; see also Wan et al., 2024). Here, we observed a similar transformational alignment, particularly in the IPS. Observing these signals in the IPS here may be due to the fact that spatial information remained relevant for the task and that the IPS and other dorsal stream areas are important for the representation of space (Mishkin et al., 1983). These findings align with the proposition that abstraction in WM is reflected in the transformation of representational structure into a future-oriented format.

Due to the sluggishness of the BOLD signal, the pre-cue delay contained a mixture of signals reflecting stimulus-processing along with WM maintenance. Consistent with past work, we show that sensory and mnemonic information is multiplexed in the IPS (Rademaker et al., 2019). Importantly, training classifiers during periods far removed from perception enabled our analyses to focus on mnemonic representations thus ruling out that the transformations observed here simply reflect a shift from perception to memory. However, training classifiers on the pre-cue delay also demonstrated a transition from retrospective to prospective (**Figure S4a**) and concrete to abstract (**Figure S4c**), albeit with greater emphasis on retrospective and concrete codes. These results indicate that the patterns observed here are robust to the specifics of the approach, but that different ways of approaching multivariate analysis can highlight more sensory-like codes on the one hand or more abstract codes on the other when both are multiplexed.

In sum, a combination of multivariate decoding, RSA, and MDS revealed that both IPS and VC initially maintained concrete past locations. Over time, these representations became progressively more abstract and prospective with abstract, prospective coding particularly prominent in the IPS. These results suggest that WM guides future cognition by reformatting sensory signals of the past into more abstract codes to guide prospective expectations.

## Acknowledgment

The authors would like to thank Michael Dicalogero for help with collecting the data. This work was supported by startup funds from Florida State University (DEN).

## SUPPLEMENT MATERIAL

**Figure S1.**
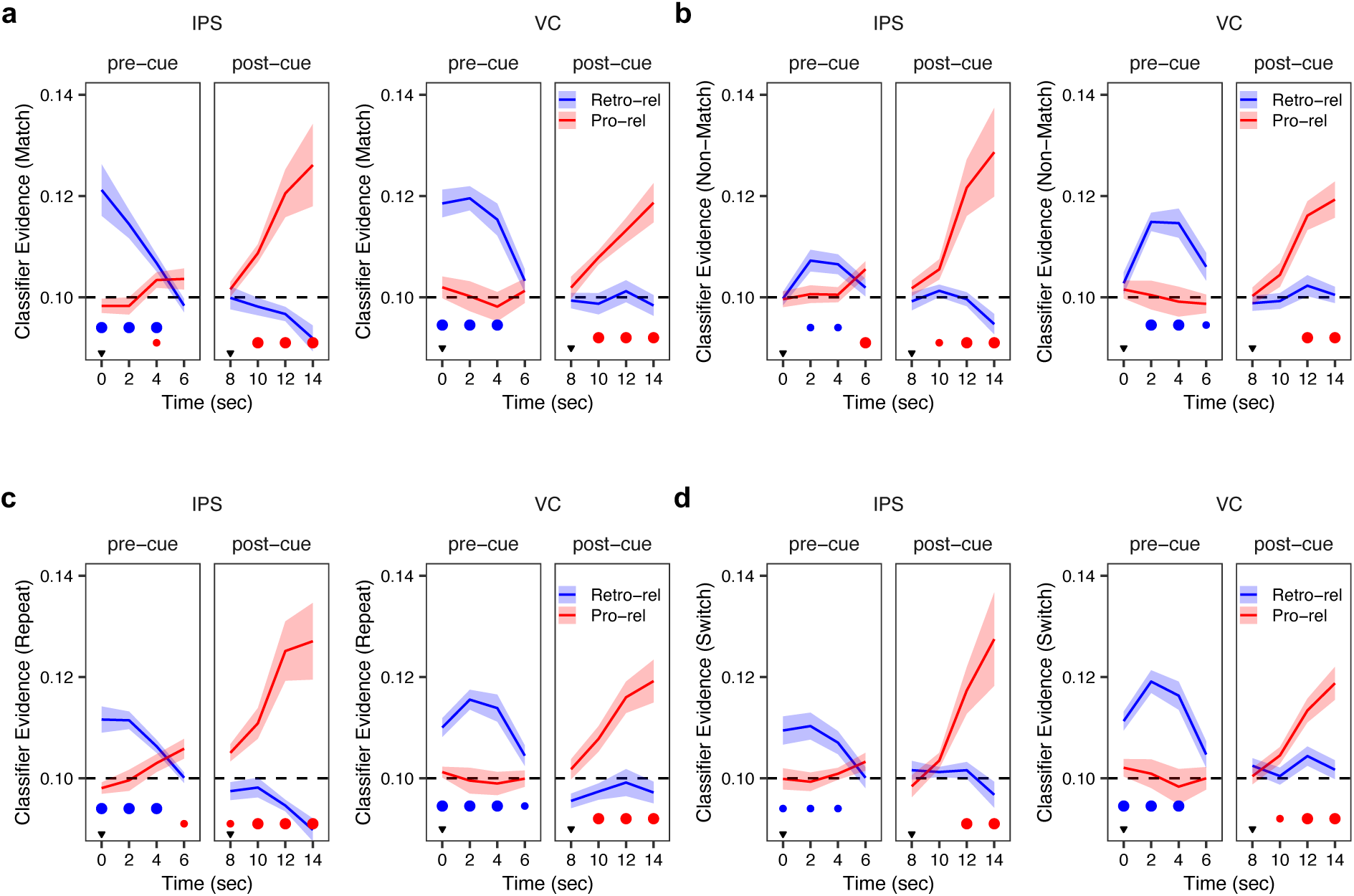
The classifier evidence of the match, non-match, repeat, and switch trials. Classifier evidence for the retrospective-relevant (‘Retro-rel’) and prospective-relevant (‘Pro-rel’) spatial locations was analyzed separately by match type of just-probed items and switch condition. The significance at each time point is denoted by small and medium dots, representing q < 0.05 and q < 0.01, respectively (FDR corrected). The inverted triangles denote the time points of the presentation of the reference stimulus and the cue, respectively. **(a)** Classifier evidence for post sequence-match trials. **(b)** Classifier evidence for post sequence-non-match trials. **(c)** Classifier evidence for repeat condition. **(d)** Classifier evidence for switch condition.

**Figure S2.**
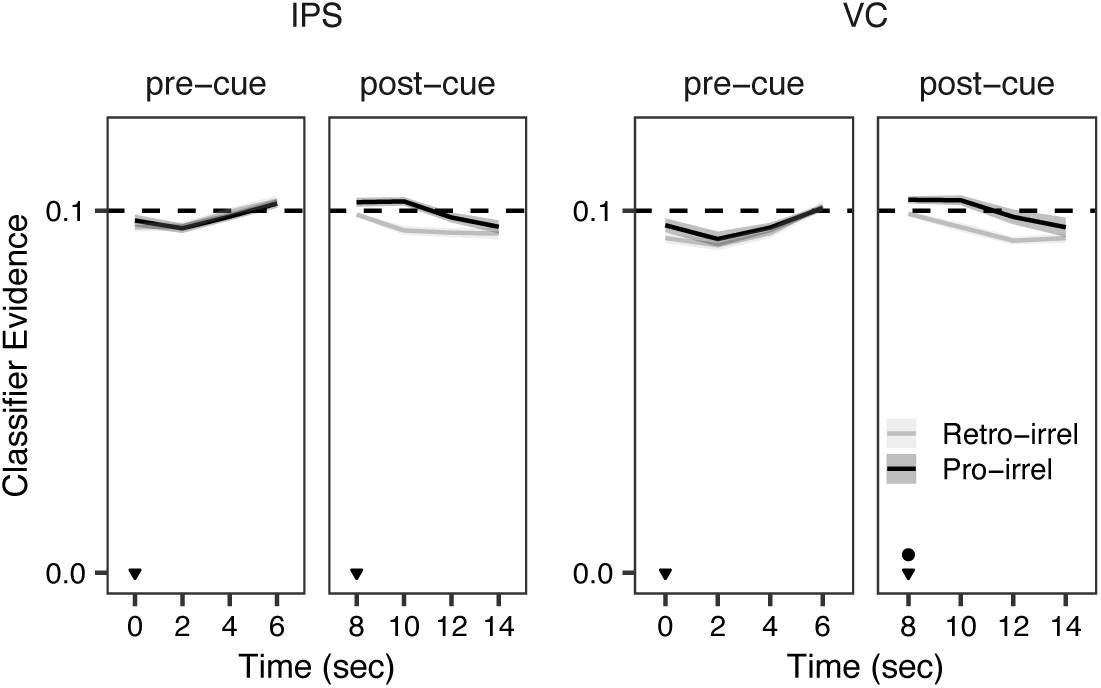
The classifier evidence for either retrospective-irrelevant or prospective-irrelevant spatial locations. Retrospective-irrelevant (‘Retro-irrel’) refers to the spatial location of the last probed item in the uncued hemifield. Prospective-irrelevant (‘Pro-irrel’) refers to the next spatial location of the last probed item in the uncued hemifield in the sequence. The significance at each time point is denoted by small dot, representing q < 0.05, respectively (FDR corrected). The inverted triangles denote the time points of the presentation of the reference stimulus and the cue, respectively. irrel (irrelevant).

**Figure S3.**
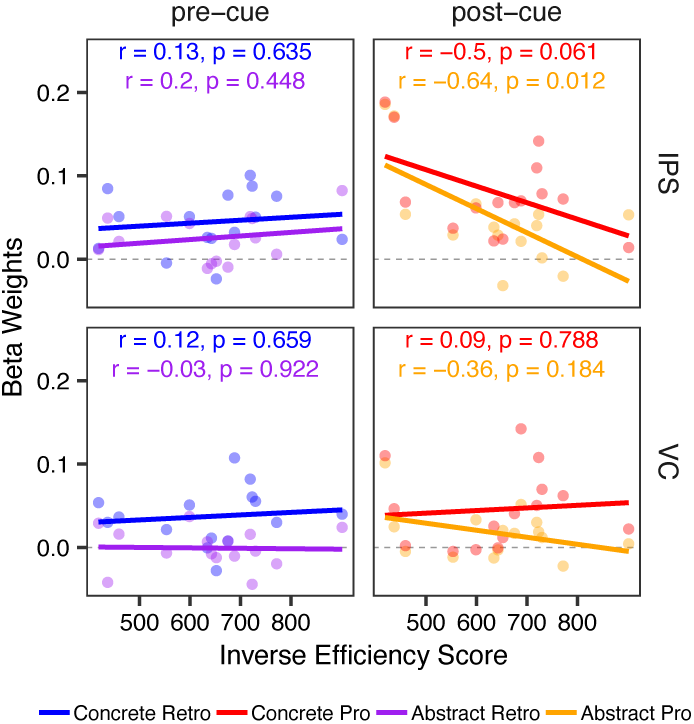
Brain-Behavior relationship. Correlation between the strength of WM codes and inverse efficiency score for each ROI. For the ‘Concrete Retro’ and ‘Abstract Retro’ conditions, beta weights were averaged over the pre-cue period (2–6 seconds). For the ‘Concrete Pro’ and ‘Abstract Pro’ conditions, beta weights were averaged over the post-cue period (10–14 seconds).

**Figure S4.**
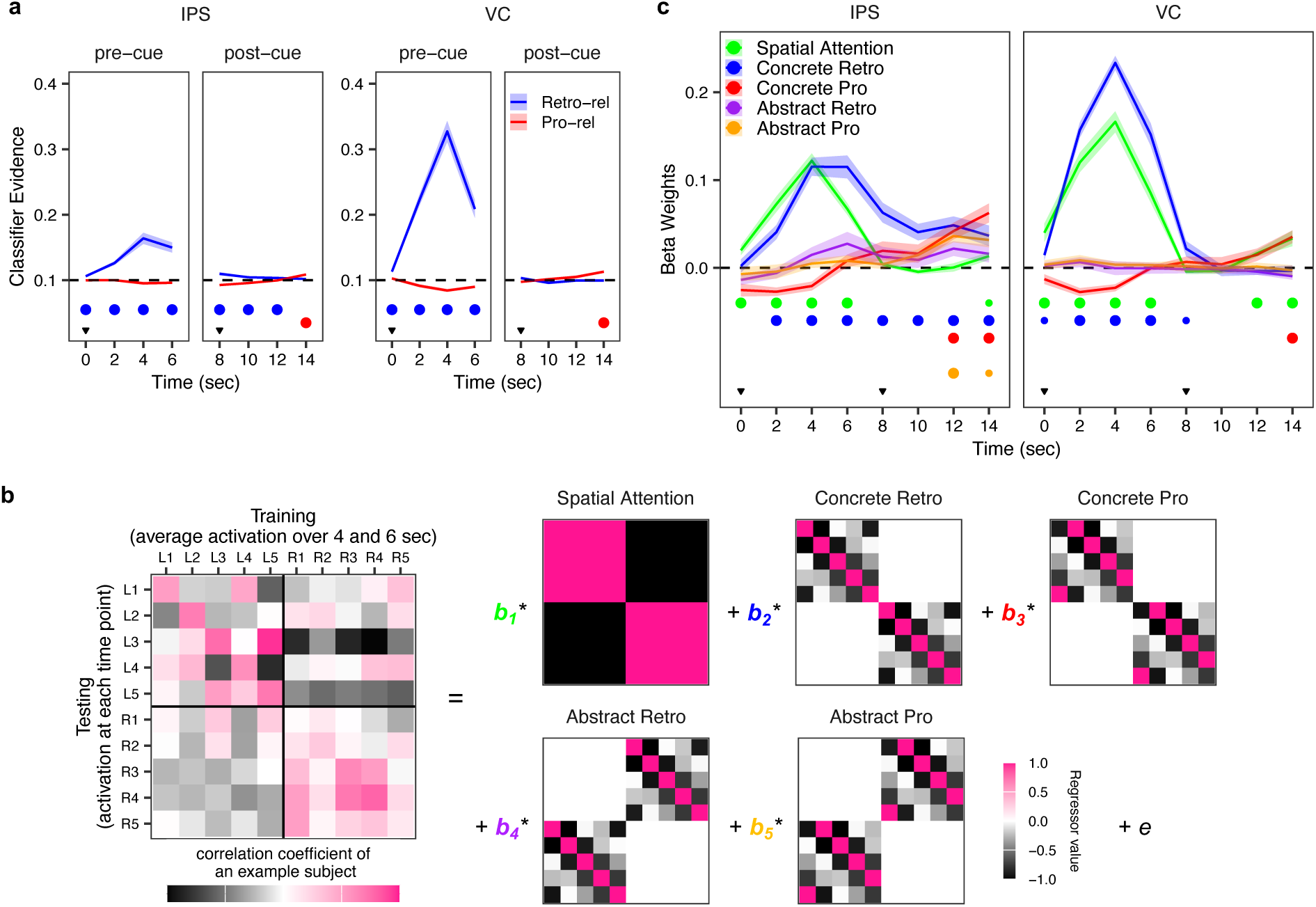
Results of the pre-cue trained classifier and RSA. In the main manuscript, we used the last two time points of the post-cue delay (12 and 14 seconds post-probe onset) as the training time period in order to focus on mnemonic representations. Here, we presented results based on the last two time points of the pre-cue delay (4 and 6 seconds post-probe onset) as the training time period. Although the sensory-related signals were prominently observed in both the temporal generalization decoding and representational similarity analyses, mnemonic-related signals (prospective and abstract codes) were also detected. **(a)** Retrospective-relevant (‘Retro-rel’) refers to the classifier evidence corresponding to the spatial location of the just-probed item. Prospective-relevant (‘Pro-rel’) refers to the classifier evidence corresponding to the next spatial location following the reference. The significance at each time point is denoted by the medium dot, representing q < 0.01 (FDR corrected). The inverted triangles denote the time points of the presentation of the probe and the cue, respectively. **(b)** The left depicts an example representational similarity matrix formed by correlating training patterns of each location with testing patterns of each location. These matrices were submitted to multiple linear regression using regressors denoting idealized matrices of spatial attention, concrete retrospective coding, concrete prospective coding, abstract retrospective coding, and abstract prospective coding. We estimated the beta weight for each regressor at each time point. Training patterns were formed by averaging the activation patterns at 4 and 6 seconds post-stimulus onset. Hypothetical models were created assuming that the training patterns reflect the retrospective location/position. Hence, retrospective models reflect a correspondence between testing patterns (e.g., L1, the first row) that correspond to the location/position preceding the training pattern (e.g., L1) with increasing dissimilarity as a function of increasing retinotopic distance. **(c)** The estimated beta weights for each regressor at each time point. The beta weights were compared against a value of 0 and corrected for multiple comparisons using false discovery rate (FDR). The significance at each time point is denoted by small and medium dots, representing q < 0.05 and q < 0.01, respectively (FDR corrected). The shaded regions represent ±1 standard error of mean. The inverted triangles in black denote the time points of the presentation of the probe stimulus and the cue, respectively.

**Figure S5.**
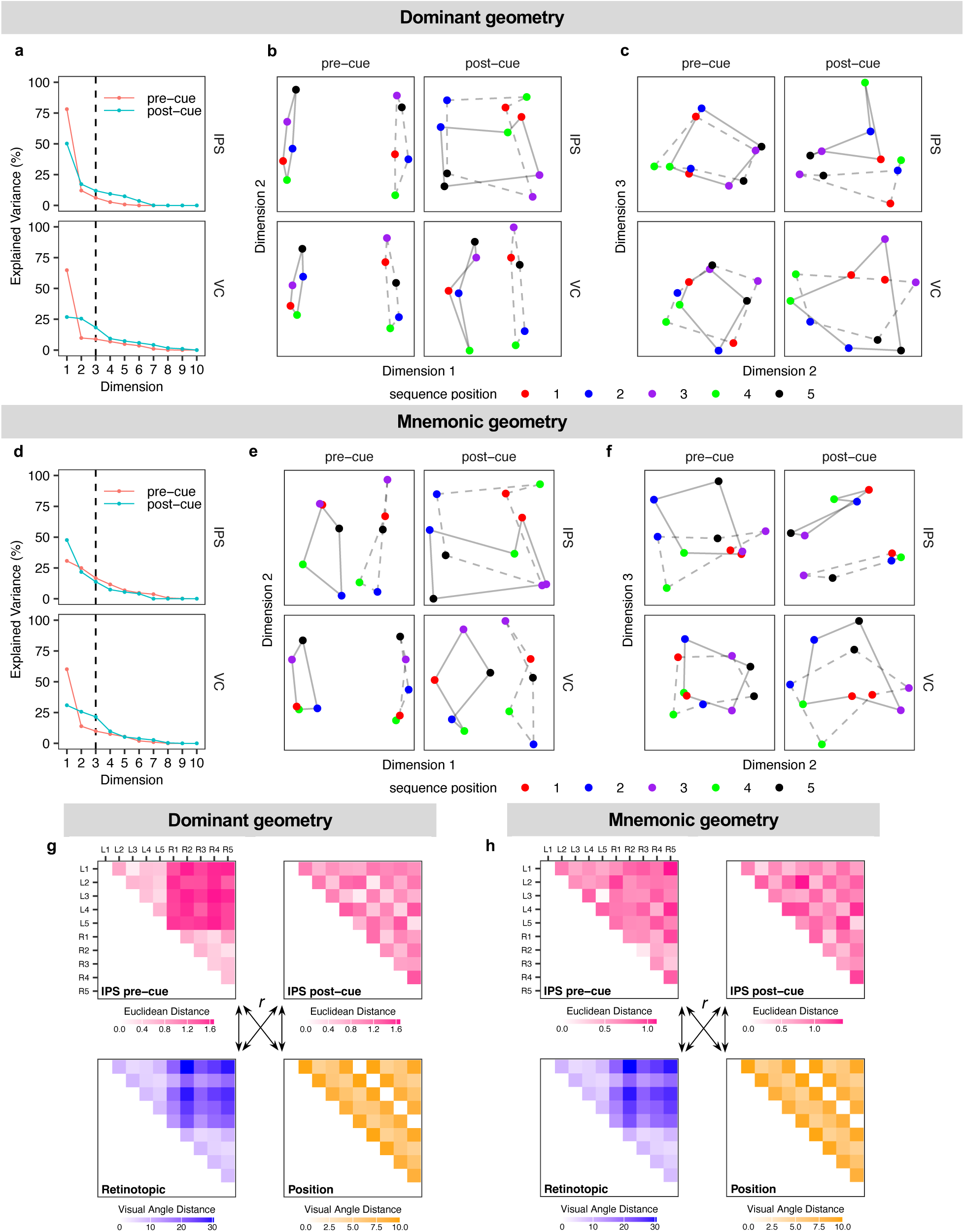
Results of the MDS analysis. **(a)** The amount of variance explained by each dimension of the dominant geometry. **(b)** WM representational space by the first and second dimensions of the dominant geometry. **(c)** WM representational space by the second and third dimensions of the dominant geometry. **(d)** The amount of variance explained by each dimension of the mnemonic geometry. **(e)** WM representational space by the first and second dimensions of the mnemonic geometry. **(f)** WM representational space by the second and third dimensions of the mnemonic geometry. **(g)** Schematic illustration of the group-level Euclidean distance in the dominant MDS geometry between spatial locations in the IPS pre- and post-cue (top row). Visual angle distance between spatial locations (bottom row). The visual angle distance was estimated considering retinotopic (bottom left) and sequence position geometries (bottom right). **(h)** Schematic illustration of the group-level Euclidean distance in the mnemonic MDS geometry between spatial locations in the IPS pre- and post-cue (top row). Visual angle distance between spatial locations (bottom row). The visual angle distance was estimated considering retinotopic (bottom left) and sequence position geometries (bottom right).

**Figure S6.**
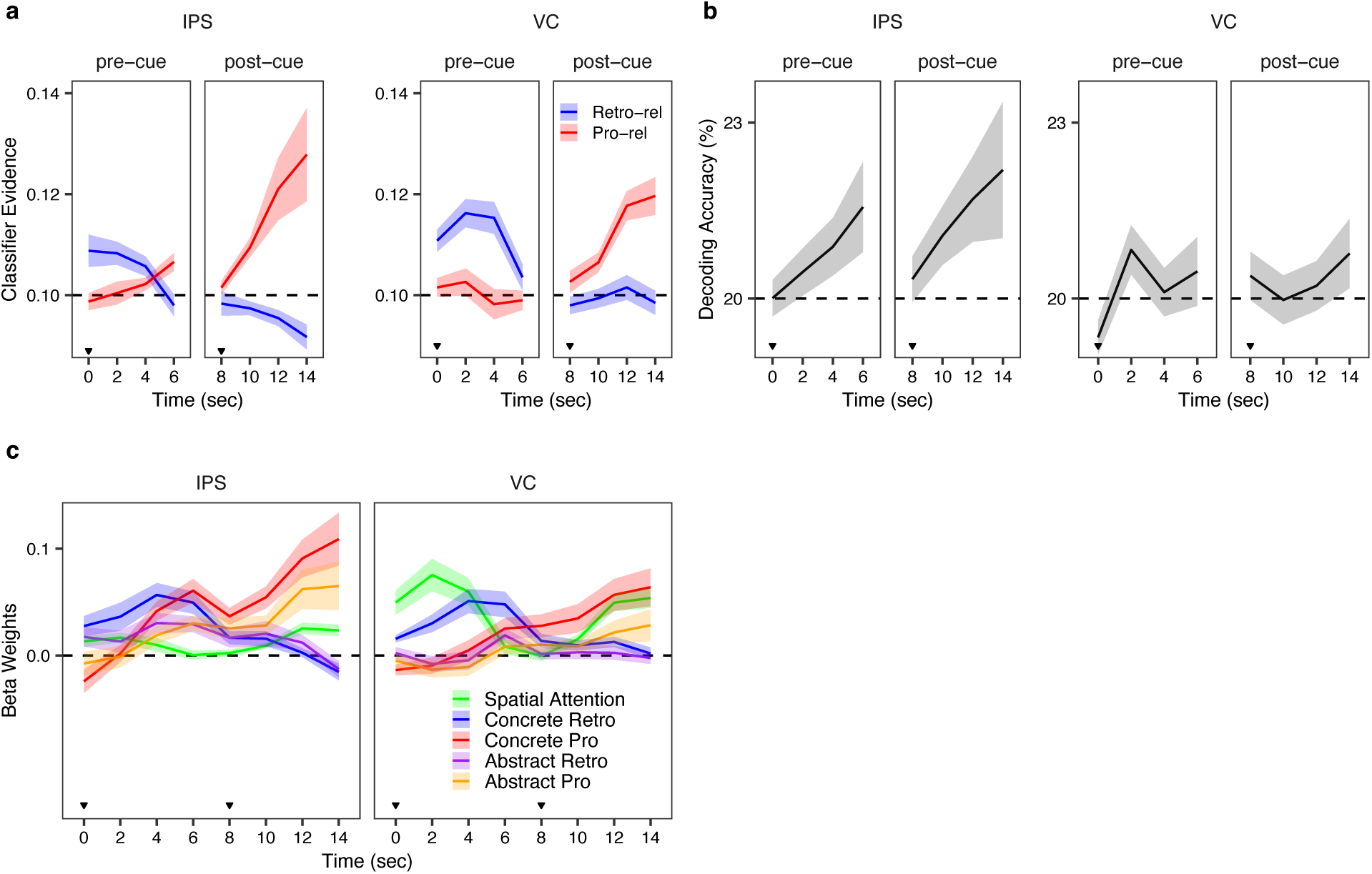
Results of MVPA and RSA analyses using the top 300 selected voxels. Analyses restricted to the top 300 voxels replicated the key patterns reported in the main manuscript. **(a)** Temporal generalization decoding analysis, replicating the pattern shown in Figure 2a. **(b)** Cross-visual hemifield decoding analysis, replicating the pattern shown in Figure 3b **(c)** Representational similarity analysis, replicating the pattern shown in Figure 5a.

